# Rapid Adaptive Increase of Amylase Gene Copy Number in Indigenous Andeans

**DOI:** 10.1101/2025.03.25.644684

**Authors:** Kendra Scheer, Luane J. B. Landau, Kelsey Jorgensen, Charikleia Karageorgiou, Lindsey Siao, Can Alkan, Angelis M. Morales-Rivera, Christopher Osborne, Obed Garcia, Laurel Pearson, Melisa Kiyamu, María Rivera-Chira, Fabiola Leon-Velarde, Frank Lee, Tom Brutsaert, Abigail W. Bigham, Omer Gokcumen

## Abstract

The salivary amylase gene (*AMY1*) exhibits remarkable copy number variation linked to dietary shifts in human evolution. While global studies highlight its structural complexity and association with starch-rich diets, localized selection patterns remain under explored. Here, we analyzed *AMY1* copy number in 3,723 individuals from 85 populations, revealing that Indigenous Peruvian Andean populations possess the highest *AMY1* copy number globally. A genome-wide analysis showed significantly higher amylase copy numbers in Peruvian Andean genomes compared to closely related populations. Further, we identified positive selection (selection coefficient of 0.0124, log likelihood ratio of 11.1543) at the nucleotide level on a haplotype harboring at least five haploid *AMY1* copies, with a Peruvian Andean-specific expansion coinciding with potato domestication (∼6–10 kya). Using ultra-long-read sequencing, we demonstrated that previously described recombination-based mutational mechanisms drive the formation of high-copy *AMY1* haplotypes observed in Andean population. Our study provides a framework for investigating structurally complex loci and their role in human dietary adaptation.

## Introduction

The amylase enzyme plays a central role in the modern human diet. Its main function is to digest starch, a cornerstone of most contemporary diets, into simple sugars ^1^. Among humans, salivary amylase (*AMY1*) copy number variation demonstrates exceptional diversity, with individuals carrying between 2 and 20 copies per diploid genome, potentially affecting salivary amylase dosage and downstream starch hydrolysis ^2^. Early duplications of *AMY1* in hominin evolution facilitated further copy number expansions in humans ^3^, which some studies have linked to agricultural diets ^2,4^. A similar trend has been observed in other species consuming broad range diets ^5^. For example, as they began cohabiting with humans, dogs experienced an increase in amylase gene copy number, especially in breeds living with agricultural communities ^5–7^. The rise in amylase copy number in dogs likely led to a higher pancreatic amylase dosage and is associated with amylase content in saliva. Further, human amylase copy number variation has been associated with metabolic traits, including the composition of the gastrointestinal microbiome, which may shape metabolic and oral health ^8,9^. Despite these exciting associations, the structural complexity of the *AMY1* locus, characterized by recurrent duplications, deletions, and inversions, has complicated efforts to connect gene dosage to local dietary evolutionary pressures in extant human populations.

The Andean Altiplano, where potato domestication began approximately 6,000–10,000 years ago ^10,11^, offers a compelling model for investigating the adaptive significance of *AMY1* copy number variation in extant human populations. Potatoes, a dietary staple in this region, are among the highest glycemic index foods, meaning that their consumption can cause spikes in blood glucose levels far more easily than other carbohydrates ^12^. For thousands of years, Andeans have relied heavily on the nutrition provided by this staple, which still constitutes over 50% of daily caloric intake in the modern Andean diet ^13–15^. Additionally, prior work has already demonstrated selection on the digestive enzyme, *MGAM*, in relation to the potato starch digestion process in Andeans ^16,17^. Thus, potatoes may have played a significant role in shaping the evolution of genes involved in carbohydrate metabolism among Andeans.

Here, we accurately genotyped *AMY1* copy numbers in a cohort of Peruvian Andean individuals, combining population genomic, haplotypic, and structural analyses. By integrating ultra-long-read sequencing with genome-wide selection tests, we reveal a signature of local adaptation affecting the *AMY1* copy number variation likely as a response to the starch-rich diet in Andean populations. These findings contribute to our understanding of local dietary adaptations and the evolutionary significance of copy number variation in complex genomic loci.

## Results

### Peruvians and Akimel O’odham/Pima have the highest ***AMY1* copy numbers among populations tested**

We calculated gene copy number variation from 3,723 individuals across 85 global populations using two read-depth-based genotyping approaches (**Table S1,** see **Methods)**. These counts were validated with Digital Droplet PCR for 101 results and 33 previously published high-quality amylase haplotypes with r^2^ values of 0.94 (p = 6.01 x 10^-11^) and 0.98 (p = 1.81 x 10^-25^), respectively ^3^. We verified previous observations that Peruvians in Lima, Peru (PEL) possess strikingly elevated *AMY1* copy numbers compared to other global populations (**Figure 1A**) ^2^. In fact, PEL display one of the highest median copy numbers (PEL median = 10, n=85) along with the Pima population (preferably known as Akimel O’odham, median = 10, n=14) among all 85 populations (worldwide median = 7) tested in this study.

**Figure 1.**
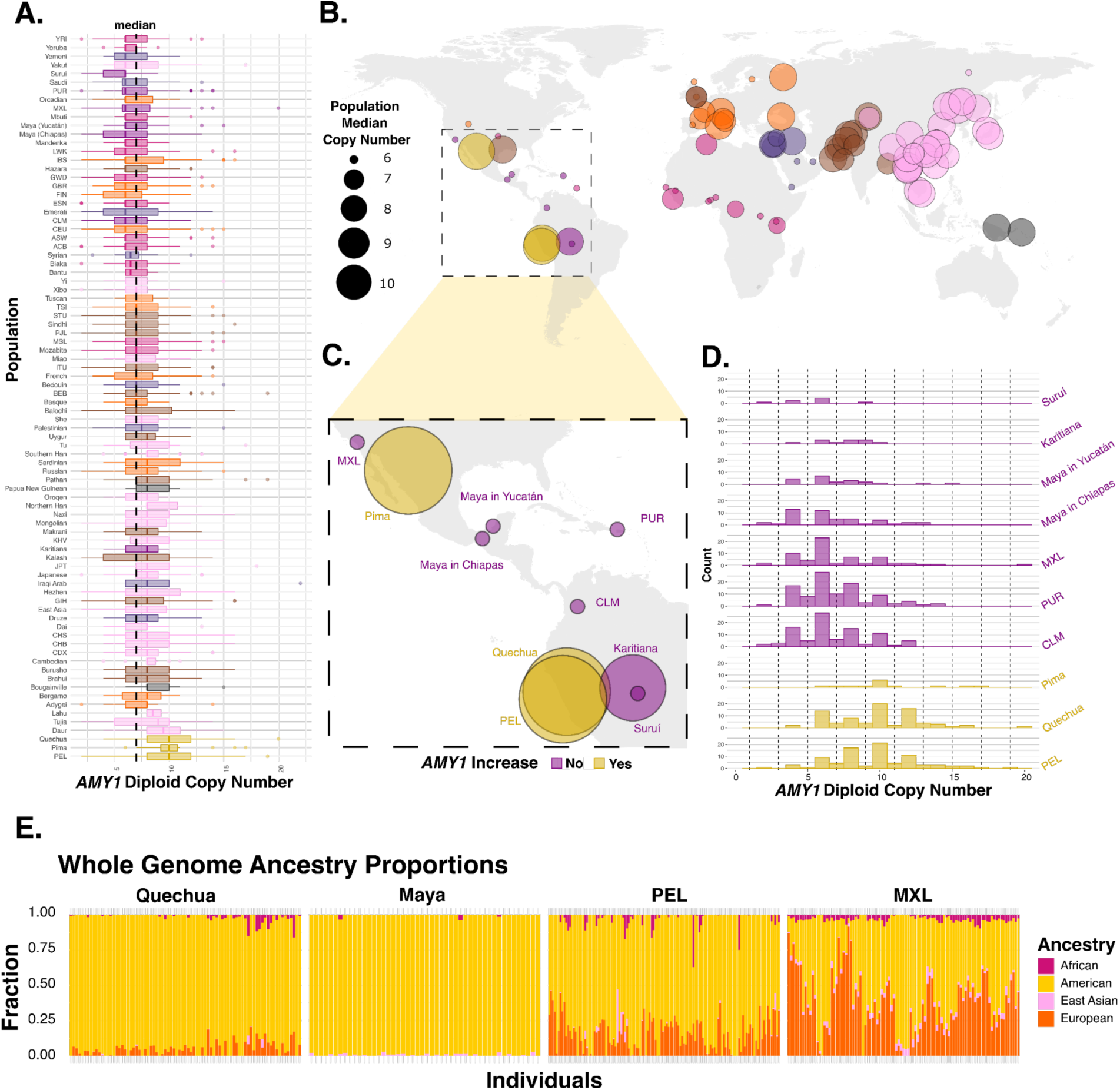
(**A**) Boxplots of *AMY1* diploid copy numbers across all populations, ranked by median. Populations are compiled using samples from the 1000 Genomes dataset, the HGDP dataset, individuals from Tibet and the Middle East, as well as Quechua and Maya individuals we have provided (see **Methods).** American populations have been split into gold (high copy number populations) and purple (low copy number populations). A black dotted line indicates the overall median (7). (**B**) Geographic distribution of all populations with circle size indicating median *AMY1* copy number (per diploid cell) and colors representing regions. Population counts are found in **Table S2** (**C**) Zoom in on American populations from panel B with populations demonstrating a higher proportion in *AMY1* genes labeled in gold. (**D**) The histogram of the *AMY1* copy numbers of the American populations from panel C, the three previously highlighted indigenous populations show an increase in the *AMY1* copy number. (E) Bar plots displaying the fraction of different ancestry proportions for populations tested in this study. The Maya from Chiapas population contains the least amount of non-American ancestry while MXL contains the most.

Prior work has demonstrated differing continent level trends regarding *AMY1* copy number ^2,18^. To assess the *AMY1* copy number pattern of variation among populations within each continent, we performed a Kruskal-Wallis (KW) test (**Figure S1A**). This analysis revealed significant population structure in the Americas (p < 2.2e-16), while other continents showed relatively homogeneous *AMY1* distributions (**Figure 1B, 1C, & 1D**). This high level of population structure is driven by PEL and Pima possessing the highest median copy numbers among all 85 populations tested, while other American populations exhibit some of the lowest median copy numbers **(Figure 1A)**. When we remove both the Peruvian and Pima populations, this population structuring is lost (KW p = 0.4) (**Figure S1B**), indicating that the observed signal is primarily driven by the high *AMY1* copy numbers in these two populations.

Due to the known European and African admixture in the American populations in the 1000 Genomes Project, we tested if this difference of *AMY1* copy number among populations can be explained by varying amounts of African and European admixture. We therefore repeated our analysis in an independent cohort of Quechua speakers from Lima and Cerro de Pasco, Peru (n = 81) and in a control cohort of Mexican Maya speakers (Tzeltal, Tzotzil, and Ch’ol) from Palenque, Chiapas, Mexico (n = 48) (**Table S1**) and likewise found exceptionally high *AMY1* copy numbers in the Quechua cohort (**Figure 1A-D**). All of these individuals have high amounts of Indigenous American ancestry (**Figure 1E**; **Tables S2-S3**). We found that the Quechua population had *AMY1* copy numbers comparable to the PEL population (both medians = 10, range = 4-20 and 2-19, respectively), while the Maya population had *AMY1* copy numbers (median = 6, range = 2-13) that did not differ significantly from other American populations from the 1000 Genomes and Human Genome Diversity Project (HGDP) datasets. We note that the individual from the PEL population that had the lowest *AMY1* copy number (2) had 100% African ancestry in the locus (**Table S4**). Of the two Quechua individuals who carried just four *AMY1* copies, one showed 100% European ancestry at the amylase locus and was therefore excluded from subsequent analyses (**Table S5**). Overall, we conclude that the observed high levels of *AMY1* copy number variation among American populations cannot be explained solely due to the percentage of Indigenous American ancestry (**Figure 1D & 1E**).

### *AMY1* copy number is a genome-wide outlier in Peruvians

We hypothesized that selection, perhaps in response to a starch-rich potato heavy diet, drove the high *AMY1* copy number in Peruvian populations. Given that the indigenous Andean population is descended from populations that have undergone several bottle necklaces (including both the crossing of the Bering Strait and the entrance to South America) which may lead to population differentiation, we conducted genome-wide analysis to rule out neutral, demographic forces. First, we conducted a genome-wide VST outlier analysis, an FST analog designed for copy number variation ^19^. We prioritized the comparison between Quechua and Maya individuals due to their relatively higher Indigenous American ancestry compared to 1000 Genomes populations (**Figure 1E**). Specifically, we compared gene copy number variants from genes that had a greater than 1 total copy number difference when rounded to an integer. We calculated these results using the program mrCaNaVaR between Peruvian Quechua individuals from Cerro de Pasco (n = 46) and Maya individuals from Palenque (n = 48) (**Figure 2A, Table S6**). Because mrCaNaVaR cannot distinguish among closely related paralogs, we applied a second copy-number caller to determine whether the observed variation occurred in *AMY1* or *AMY2*. This analysis confirmed that the increase was specific to *AMY1*, as *AMY2A* and *AMY2B* showed no significant variation across American populations (**Figures S2** & **S3**). *AMY1* ranked among the top three most differentiated genes genome-wide in copy number, placing in the top ∼0.07% of all genes tested (**Table S7**).

**Figure 2:**
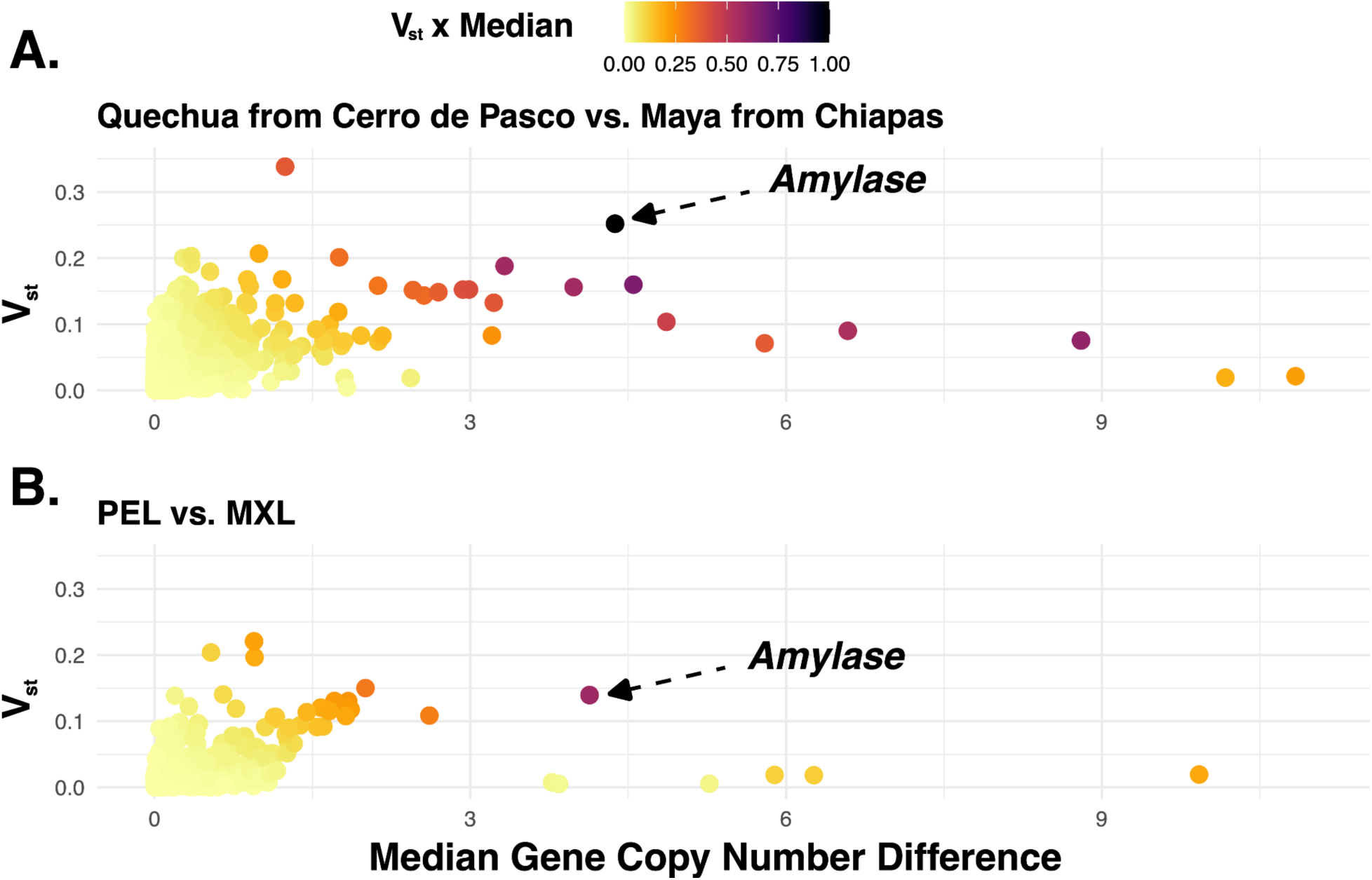
(**A**) Scatterplot showing the relationship between Vst and median gene copy number difference in all protein-coding genes in Quechua from Cerro de Pasco and Maya from Palenque populations for genes that have a total copy number difference greater than 1. Genes that have zero difference between the populations (both Vst and the Median difference = 0) are not plotted. The *AMY1* and *AMY2*, are represented by a single dot that has been pointed out. Darker shading is associated with greater Vst and median copy number difference values. (**B**) Scatterplot showing the relationship between Vst and median gene copy number difference in all protein-coding genes in Peruvians in Lima and Mexicans in Los Angeles populations for genes that have a total copy number difference greater than 1.

To further validate our findings, we replicated this analysis using 1000 Genomes data, comparing PEL and Mexicans from Los Angeles (MXL) (**Figure 2B, Table S6**). This analysis corroborated our results, with *AMY1* copy number variation ranking among the highest median differences observed for common copy number variable genes in the top seven genes (top ∼1.2% **Table S7**). The slight differences in the two comparisons may be partially influenced by increased admixture in the PEL and MXL populations. Nevertheless, population differentiation tests across two independent cohorts consistently showed that variation in *AMY1* copy number exceeds genome-wide expectations, encouraging us to further investigate the hypothesis that high copy numbers in the Indigenous Andean population may result from a selective sweep.

### Identifying haplotypes harboring high *AMY1* copy numbers in Peruvian Andean populations

To understand the haplotype structure associated with high *AMY1* copy numbers, we analyzed linkage disequilibrium (LD) using the Kruskal-Wallis test at the amylase locus and its flanking regions (hg38 Chr1:103,348,464–103,830,994) across populations in the 1000 Genomes Project and compared these patterns to those observed in the Quechua and Maya populations (**Figure S4; Table S8**). We found that in most populations, flanking single nucleotide variants (SNVs) were not significantly correlated with *AMY1* copy number. However, we identified 81 SNVs that showed significant correlation with *AMY1* copy number and were shared between East Asian and Andean/Maya populations, suggesting the presence of a shared ancestral haplotype (**Table S9**). These SNVs also were in long-range, high LD (r^2^) with each other in the Quechua population (**Figure S5, see Github**).

Further, we identified 7 region-specific SNVs that appear to have evolved on this ancestral haplotype in Indigenous American populations as well as 17 SNVs private to the Andean populations (**Table S9**). These variants were also in strong LD with one another and weaker LD with the ancestral haplotype due to their later addition (**Figure S5**). The combination of these SNVs form a population-specific haplotype block in Andeans that extends further on each side of the amylase locus than any other observed among the populations analyzed, and this haplotype was significantly correlated with high *AMY1* copy numbers (**Figure S4**).

We hypothesized that if the increased LD at the amylase locus in the indigenous Andeans reflects a recent selective sweep, then the same SNVs that correlate with *AMY1* copy number should exhibit increased FST between Peruvian Quechua and Maya populations, mirroring the trends observed in our VST outlier analysis. Consistent with these expectations, we found that the SNVs tagging the *AMY1* copy numbers show significantly high FST values (ex. rs58677276 = 0.311 and rs143597860 = 0.199 are both above 99^th^ percentile) compared to the rest of chromosome 1 (**Figure 3A**, **see Github**).

**Figure 3:**
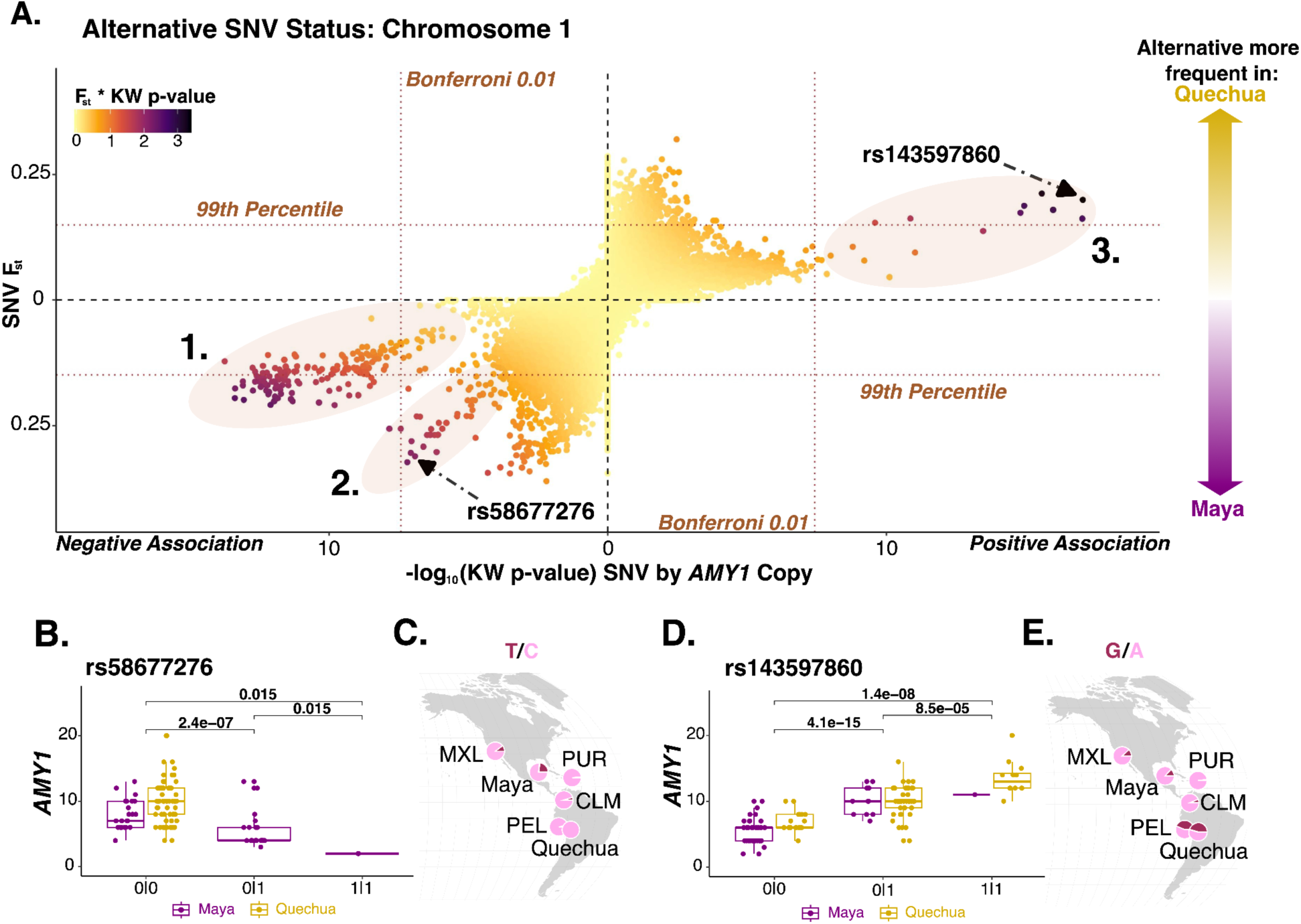
(**A**) Chromosome 1 SNVs in Quechua from Peru (top) and Maya from Chiapas (bottom) with Indigenous American ancestry in the amylase region (**Table S4**) in the alternative allele. Y-axis: directional FST with SNVs that have a higher frequency in the Quechua population above the dotted axis and SNVs with a higher frequency in the Maya population below the dotted y-axis. The 99th percentile threshold is marked. X-axis: -log10(p-values) from a Kruskal-Wallis test comparing diploid *AMY1* copy numbers among genotype classes. SNVs to the right of the axis are associated with an increase in *AMY1* copy number and those to the left are associated with a decrease in *AMY1* copy number. The vertical dashed line marks the Bonferroni-corrected significance threshold (α = 0.01); points further from the origin are significant after correction. Colors indicate the relationship between these statistics. Three groups of SNVs that seem to be associated with *AMY1* copy number are highlighted. (**B**) rs58677276 which we have highlighted as the example SNV for group 2. Pairwise Wilcoxon values are labeled. All SNVs in this group are negatively associated with the *AMY1* copy number and primarily appear in the Maya population. (**C**) Frequency of rs58677276 in American populations. (**D**) rs143597860 which we have highlighted as the example SNV for group 3. Pairwise Wilcoxon values are labeled. SNVs in this group are associated with an increase in *AMY1* copy number. (**E**) rs143597860 frequency in American populations.

When jointly considering population differentiation and correlation with *AMY1* copy number, we identified three SNV clusters corresponding to distinct haplotypes that drive the observed differentiation between the Maya and Quechua at the amylase locus (**Figure 3A**). The first group comprises SNVs where the reference allele was associated with higher *AMY1* copy number. The reference allele of an example SNV from this cluster, rs1930184, segregates at ∼74% in the Quechua and is also common in East Asian populations (∼60-68%), likely contributing to the higher *AMY1* copy number observed in both groups. In Maya, the reference allele was found in only ∼42% of the population, contributing to the high FST. The SNVs in this cluster were the same aforementioned SNVs that share high LD between the EAS and Andean/Maya populations. A phylogeny of the SNVs in the flanking regions of the amylase locus displayed a high copy number clade that can be tagged by rs1930184 within the East Asian populations in which most of the high copy number haplotypes in Andeans and Maya were all nested (**Figure S6**).

The second group, here tagged by rs58677276, contains SNVs where the alternative allele was associated with lower *AMY1* copy number (**Figure 3B**). The alternative allele for rs58677276 was present at frequency of ∼26% in the Maya and completely absent in Quechua (**Figure 3C**) though other SNVs in this cluster were present at frequencies of ∼1-2% in Quechua. The third cluster, tagged here by rs143597860, was specific to the Americas and includes SNVs where the alternative alleles were associated with high diploid *AMY1* copy numbers (**Figure 3D**). Rs143597860 segregated in its alternative state at ∼46% in Quechua but only at ∼14% in Maya (**Figure 3E**) and had the highest Pearson correlation coefficient value between a SNV and *AMY1* copy number (Pearson correlation 0.71, p-value = 3.3×10^-13^) (**Table S10**).

We hypothesize that this population-specific haplotype, tagged by rs143597860 and harboring five or more *AMY1* copies, provides evidence that its frequency has increased in Indigenous Andean populations through a soft selective sweep.

### Selection on standing variation best explains the high allele frequency of high *AMY1* copy number harboring haplotypes in indigenous Andeans

To assess whether demographic bottlenecks alone could account for the observed allele frequency differentiation of high *AMY1* copy number haplotypes, we conducted neutral simulations using msprime (see **Methods**). Using published divergence times and effective population sizes, along with our own estimates via Relate (**Figure S7**), we were unable to replicate the observed allele frequency of rs143597860 (**Table S11**). Specifically, across 3×10^9^ simulated base pairs (roughly the size of the human genome) we could not replicate a scenario where any mutated SNPs achieved a frequency of greater than 45% in Andeans and less than 13% in Maya under the estimated parameters. To further test whether higher genetic drift could account for the observed frequencies, we performed 192 additional simulations under progressively reduced (and increasingly unrealistic) effective population sizes for the Quechua and Maya populations (**Figure S8**, **Table S11**). Only under highly specific and implausible conditions, such as a Quechua Ne of 250 and a Maya starting Ne of 1,000 with a final Ne of 500, did the simulations yield allele frequencies approaching the empirical data. These findings suggest that drift alone is unlikely to explain the elevated frequency of this allele in Indigenous Andeans, pointing instead to a potential role for positive selection.

Our observation that SNVs that are in unusually high correlation with *AMY1* copy numbers exhibit significant population differentiation in Quechua suggests that LD patterns for these SNVs should differ markedly between Andean Peruvians and the closely related Maya population. To test this empirically, we calculated cross-population extended haplotype homozygosity (XP-EHH) between Quechua and Maya genomes ^20^. As a sanity check for the accuracy of this test, we verified that the *EGLN1* locus, a known target of high-altitude selection in Andean populations ^21^, also displayed significantly high XP-EHH values (>4, >99^th^ percentile, **Figure S9, see Github)**, replicating previous studies and supporting the sensitivity of our analysis. We found that multiple SNVs flanking the *AMY1* locus, including SNV rs143597860, also displayed chromosome-wide significant XP-EHH values more significant that what was observed for *EGNL1* locus (>6, >99^th^ percentile **Figure 4A**, **Figure S9, see Github**). This result is conservative, as most genetic maps are based on SNV data and exclude recombination events within the *AMY1* locus due to segmental duplications that evolve more rapidly than SNVs. Moreover, the significant XP-EHH block flanks the *AMY1* locus on both sides, containing multiple linked SNVs associated with *AMY1* copy number.

**Figure 4:**
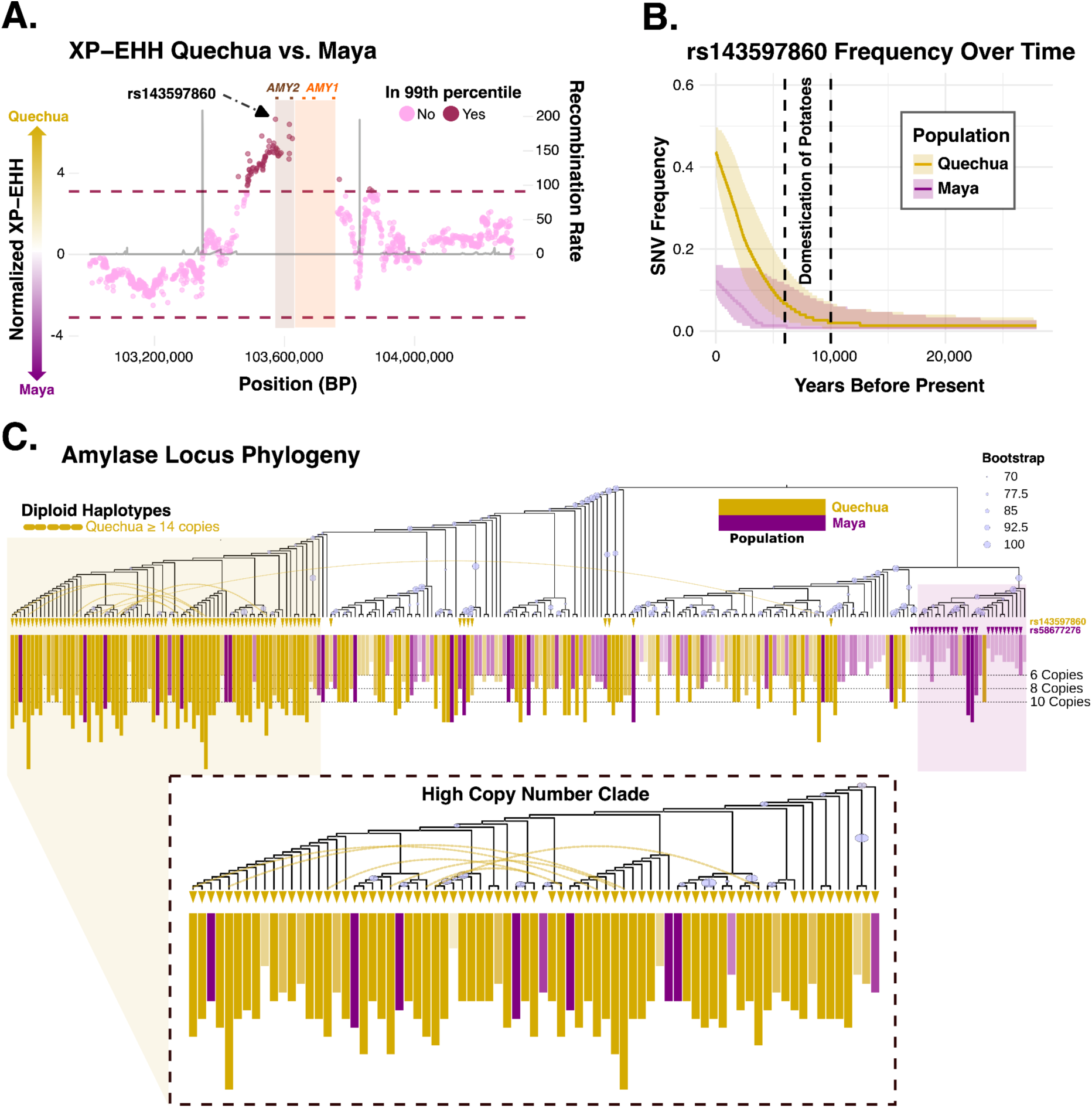
(**A**) XP-EHH between Quechua and Maya samples in the amylase locus and flanking regions along with regional recombination rates. Brown squares are *AMY2B* and *AMY2A* while orange squares are *AMY1A*, *AMY1B*, and *AMY1C*. Chromosome-wide significant values are in red. Positive XP-EHH values have longer haplotypes in Quechua; negative XP-EHH values have longer haplotypes in Maya. (**B**) Temporal trajectories of the derived allele rs143597860 in Quechua and Maya populations. Solid lines trace the maximum-a-posteriori (MAP) estimate of derived-allele frequency through time for each population. Shaded regions mark every frequency bin with non-zero posterior probability at each generation (i.e., the full posterior support and not a fixed-probability credible band). Dashed vertical bars denote the archaeologically inferred interval for potato domestication (6,000–10,000 BP). (**C**) A phylogeny of the haploid amylase and flanking regions. Bars are colored by population and shaded by *AMY1* copy number, with the presence of rs58677276 **(Figure 3B, C)** and rs143597860 **(Figure 3D, E)** labeled. All Quechua individuals with ≥ 14 *AMY1* copies (exceeding the max Maya copy numbers) have shared haplotypes linked by a dotted line.

To further evaluate the putative soft selective sweep, we conducted additional tests to ensure that our observations were not confounded by admixture or drift. First, we applied the Population Branch Statistic (PBS) ^22^, which builds on FST by incorporating an outgroup and is particularly useful for detecting selection in admixed populations. Using the FIN population as an outgroup and comparing PEL and MXL populations, we found that the tag SNV rs143597860 had one of the highest PBS values among all SNVs analyzed (top 0.04%), placing it well above the 99^th^ percentile (**Figure S10, see Github**). In parallel, we used Ohana ^23^, which explicitly models admixture components across the genome and identifies SNPs with allele frequencies that deviate from genome-wide expectations. Running Ohana on the Quechua genomes using the Iberians in Spain, (IBS) and Yoruba in Ibadan, Nigeria (YRI) populations as comparisons, we found that rs143597860 again ranked in the top 0.01% of SNVs analyzed (**Figure S10, see Github**). Together, these selection scans consistently reveal elevated population differentiation and extended haplotype homozygosity at rs143597860, providing robust, multi-method evidence for recent positive selection on this haplotype in Peruvian Andeans.

### The putative soft sweep coincides with potato domestication in the Andes

The Atlas of Variant Age with a 25-year generation time, dates rs143597860 as 7,068 years old based on a recombination clock, 27,368 years old based on a mutation clock, and 17,945 years old with a joint clock estimate, coinciding with early migrations to the Americas around 13,000-23,000 years ago ^24,25^, based on 1000 Genomes and SGDP (Simons Genome Diversity Project) (**Figure S11**)^26–28^ . As this SNV segregates at a frequency of 3.7% in populations from Central Asia Siberia, rs143597860 likely existed among the ancestral population to both Maya and Quechua, increasing in frequency among Andeans in a manner concordant with selection on standing variation due to the hypothesized selective sweep on the high amylase copy numbers.

To explore the evolutionary trajectory of rs143597860 and the likelihood of selection, we leveraged Relate to infer approximate ancestral recombination graphs and CLUES to estimate allele frequency trajectories over time, including the strength and likelihood of selection ^29,30^. These analyses revealed that the allele G, the alternate allele of the putatively selected SNV rs143597860, segregated at very low frequencies (0.7-2.7%) in ancestral populations prior to the divergence of Quechua and Maya populations, before rising in frequency after ∼10,000 years ago (**Figure 4B & S12, see Github**). In the Maya, the allele rose modestly, slightly exceeding 10% frequency in the last 10,000 years. However, the low log-likelihood ratio (LLR) score (1.53) was statistically indistinguishable from neutrality and any evidence for selection on this locus in the Maya was inconclusive. In contrast, the rs143597860 trajectory in the Quechua population showed a striking increase from ∼2% to ∼45%, best explained by a selection coefficient of 0.0124 with a log-likelihood ratio of 11.15. This result strongly supports the action of a selective sweep. Notably, the timing of this allele frequency increase coincides with the period of potato domestication in the Andean region (6,000–10,000 BP), raising the intriguing possibility that dietary shifts may have contributed to the selective pressures acting on this locus. For comparison, the *EGLN1* variant rs1769792, previously identified as a top target of selection at high altitude ^31,32^ has a selection coefficient (S) of 0.005 and an LLR score of 8.08 (**Figure S13**).

To put the aforementioned SNVs in context and to visualize the amylase haplotypic variation, we constructed a phylogenetic tree using phased SNVs from Quechua and Maya individuals encompassing the amylase locus and flanking regions (hg38: Chr1: 103,348,464-103,830,994) (**Figure 4C, see Github**). On this phylogeny, we superimposed the diploid *AMY1* copy number and the presence or absence of the three high FST SNVs described above. We found that rs58677276 appeared mainly in a clade (corresponding to the 2^nd^ cluster in **Figure 3A**) harboring diverse haplotypes that were predominantly composed of Maya individuals (25/27 haploid genomes) with low *AMY1* copy numbers (median = 4). This phenomenon may have been caused by a deletion event triggering the loss of *AMY1A* and *1B*, the core mutational unit described previously ^3,18^, and resulting in a static single *AMY1* copy number, which may be present in this clade’s haploid genomes.

More relevant to the proposed selective sweep, we identified a clade that predominantly carried the alternative allele for rs143597860 (corresponding to the 3^rd^ cluster in **Figure 3A** and the highlighted high copy number clade in **Figure 4C**), associated with the selected haplotype. This clade was largely Quechua (67/77 haploid genomes) and exhibited high *AMY1* copy numbers (median = 12), accounting for over half (64 of 116) of all haploid genomes in our dataset who have diploid *AMY1* copy numbers ≥ 10. When expanding the phylogeny to include EAS populations, we found this clade was nested within the high copy number clade tagged by rs1930184 (**Figure S6**).

### Structural haplotypes of Peruvian populations

The structural haplotypic variation of the amylase locus in Peruvians is not well-characterized. Previous studies showed that *AMY1* copy number variation involves non-allelic homologous recombination (NAHR) of a mutational unit comprising *AMY1A* and inverted *AMY1B* (**Figure S14**) ^3^. To address whether the unusually high *AMY1* copy number haplotypes in Peruvian Andeans were due to the same mutational mechanism, we analyzed data from 40 existing haplotype assemblies, including 10 Peruvian samples (**Figure 5A**; **Table S12**) ^2,3^. The existing assemblies allowed us to construct a pangenome of the amylase locus representing 33 distinct structural haplotypes (**Table S12**, **see Github**). Among these, 14 haplotypes were present in Peruvians, yet all of them also occurred in other global populations (**Figure 5B**).

**Figure 5:**
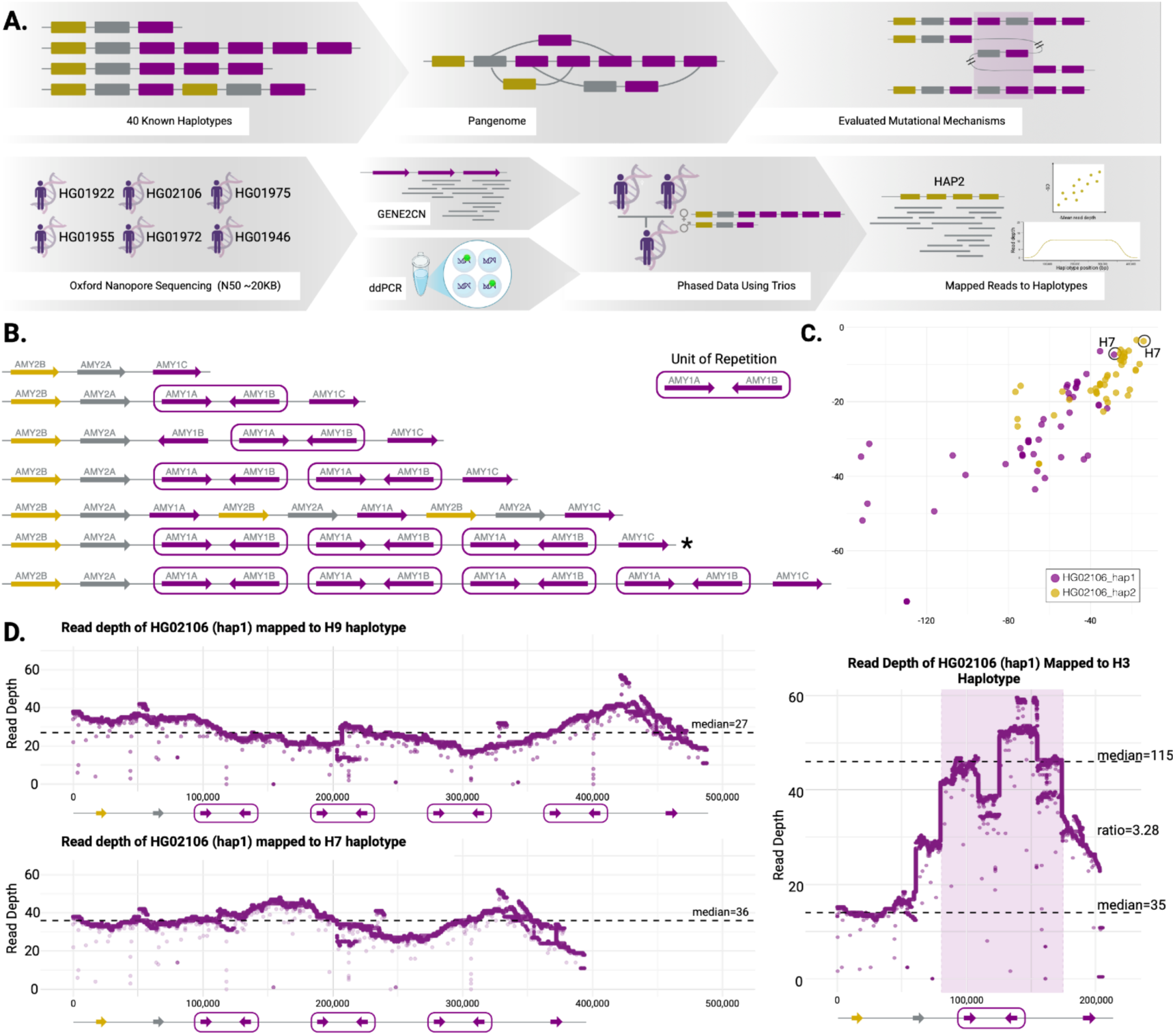
Peruvian populations have uniquely long haplotypes while maintaining the same duplication mechanisms. (**A**) Overview of the analytical approach used to identify and characterize Peruvian haplotypes and their underlying mutational mechanisms. (**B**) Haplotypes found in Peruvian individuals. Known haplotype structures reported in previous studies are shown alongside newly inferred Peruvian haplotypes identified in this study. The asterisk corresponds to the manually curated H7 haplotype present in Peruvians. (**C**) Scatter plot illustrating the relationship between mean read depth (y-axis) and standard deviation (x-axis) for individual HG02106’s reads mapped against all known haplotypes. The highlighted data points represent the best-fitting mappings for HG02106’s two haplotypes (H7), characterized by their lower standard deviation values. (**D**) The upper panel on the left shows the per-nucleotide read depth for HG02106 when aligned to the H9 haplotype. In contrast, the lower panel shows the alignment to the inferred H7 haplotype, demonstrating higher median read depth and reduced standard deviation, suggesting a better haplotype fit. On the right, the read depth when aligned to an H3 haplotype, shows the ratio between the region with invariable copy number of *AMY2B* and the region comprising the repeat *AMY1A*/*AMY1B*.

To further investigate, we generated long-read sequencing data from six PEL individuals representing a range of copy number alleles, including those carrying the highest *AMY1* copy numbers **Figure 5A**). These six PEL samples were children in independent trios with previously sequenced genomes (**Supplementary Methods)**. Specifically, we generated Oxford Nanopore long-read sequencing data ranging from 14–45× coverage (**Supplementary Table S13**), achieving an N50 of ∼20 kb. For two samples, we additionally obtained ultra-long reads with an N50 of ∼50 kb. Using this dataset, we constructed phased contigs through local assembly tools (**Figure S14, Supplementary Methods, Table S13**). We analyzed the pangenome and individual contigs for sequence signatures indicative of previously characterized microhomology-mediated break-induced replication (MMBIR) events, which have been implicated in rare occurrences of amylase duplications ^3^. No such signatures were detected in the Peruvian samples, either within the pangenome or the contigs (**Supplementary Methods**), suggesting that observed structural variation at the *AMY1* locus was not driven by MMBIR mechanisms in this population. This conclusion is further supported by the absence of significant variation in *AMY2A* and *AMY2B* copy numbers in Peruvian Andeans (**Figure S2 & S3**), which are shown to be typically affected by MMBIR.

Next, we examined the above-mentioned 40 assembled haplotypes to confirm that the observed *AMY1* copy number variation in the Peruvian samples could be explained by the recently described non-allelic homologous recombination (NAHR) mechanisms involving both *AMY1A* and *AMY1B* (**Figure 5B**). We competitively mapped the newly sequenced phased long reads from six PEL individuals (12 phased samples) onto the 40 assembled haplotypes. For each phased sample, we calculated the mean read depth and standard deviation of read depth across all assembled haplotypes to identify the best fit (e.g., **Figure 5C, Supplementary Methods**). We assumed that the best-matching haplotype would exhibit the most uniform coverage across assemblies (i.e., lowest standard deviation) and an appropriate mean read depth (see **Supplementary Methods**). These assignments were further validated by copy number estimates derived from GeneToCN and Droplet Digital PCR analyses (**Table S1)** ^3^. Overall, we successfully assigned 8 out of 12 newly sequenced phased samples to known reference haplotypes (**Table S13, S15**).

Two phased samples (HG01975 and HG02106), with an expected 14 diploid copies each, did not match the expected copy numbers by the read-depth approach (**Supplementary Methods**). We manually inspected the mapping to known haplotypes and found that the alignments to other known 3-, 5-, and 9-copy haplotypes were not uniform. Furthermore, despite being the best match, we observed apparent gaps in the mapping even for the 9-copy previously resolved Peruvian haplotype (**Figure 5D**). This led us to hypothesize that, despite structural similarities, the Peruvian haplotypes may differ in local sequence variation. To test this, we generated an estimated 7-copy haplotype by deleting one mutational unit from the 9-copy Peruvian haplotype (**Figure S16**). This adjustment significantly improved the mapping (**Table S15**), providing the best fit for these phased samples (**Figure 5C**), and allowed us to conclude that this individual may carry two 7-copy haplotypes (**Figure 5D**), and observed exclusively, with the exception of two African individuals, in East Asian and American populations to date ^33^.

Together, these results suggest that the majority of high *AMY1* copy number haplotypes observed in Peruvians already existed in global populations and have evolved through the same mutational mechanisms reported earlier ^3^. Most Peruvian haplotypes have been documented previously, except a seven-copy haplotype, which is within the expected realm of mutational possibilities but may harbor Peruvian-specific variants at the local scale.

## Discussion

The amylase locus has long been discussed in the context of human dietary adaptation. Previous studies show that the salivary amylase gene (*AMY1)* exhibits remarkable copy number variation, evolving rapidly and showing associations with dietary preferences among human populations ^2–4^. However, direct evidence of local selection on *AMY1* has remained elusive. Here, we provide robust evidence for recent selection acting on *AMY1* copy number in Peruvian Andeans. Furthermore, we show that previously described recombination-based structural mutations, specifically those involving duplications or deletions of two *AMY1* genes in a single mutational step, drive the formation of extremely high-copy *AMY1* haplotypes in Peruvians. Our findings have broad implications for understanding the genetic basis of dietary adaptations to local ecologies and the role of selection in shaping structurally complex loci.

Previous studies have shown that *AMY1* copy number increased in the human lineage prior to agriculture ^34^ and has been shaped by frequent gains and losses. This same increase in amylase copy number has been demonstrated in species consuming a broad range diet that have cohabitated with humans after the agricultural revolution ^5–7^. Thus, high-copy *AMY1* haplotypes likely existed in populations ancestral to Andean Peruvians. SNV analyses on the flanking regions of the amylase locus indicate that the high-copy *AMY1* haplotypes found in modern day Andean Peruvians are shared with high-copy haplotypes found in East Asians. Our results suggest that a soft selective sweep dramatically increased the allele frequency of these high-copy haplotypes in the Andean Peruvian population. This observation aligns with recent work on human adaptive evolution emphasizing the role of local selection on standing variation ^35^, as demonstrated convincingly for skin color ^36,37^.

It also is noteworthy that *AMY1* copy number variation emerges from recurrent NAHR events. The rates of these events are likely influenced by haplotype length where a higher number of *AMY1* gene copies lead to a higher probability of NAHR. The selective sweep that led to higher allele frequencies of high-copy haplotypes may lead to additional *de novo AMY1* copy number variation. This *de novo* variation could have been subject to selection alongside pre-existing variation. Therefore, it is plausible that both selection on standing variation and *de novo* mutations contributed to the elevated frequency of amylase haplotypes with five or more haploid *AMY1* copies. This could have led to an increased proportion of Andean individuals with high-copy haplotype. An open question remains as to whether there is an upper limit to *AMY1* copy number expansion and variation. Potential constraints may arise from genomic instability as a function of increased repeated segments or homologous alignment issues caused by copy number mismatches between maternal and paternal chromosomes. Investigating these limitations presents an exciting avenue for future research.

Our observation that *AMY1* copy number has adaptively increased in the Andean Peruvian population raises the broader question of the adaptive function of salivary amylase. At the molecular level, the enzyme’s primary role is starch digestion ^38,39^, and there is accumulating evidence that increased *AMY1* copy number leads to a higher dosage of functional paralogs, resulting in elevated salivary amylase levels ^8,40^. However, how this increased enzyme dosage translates into organismal-level function remains unclear. Several studies have reported associations between higher *AMY1* copy number and shifts in the oral and gut microbiomes. For instance, elevated copy number has been linked to greater oral microbiome diversity ^8^ including increased abundance of pathogenic *Streptococcus* species in the oral cavity, which may underlie the observed association between high *AMY1* copy number and denture use in a large UK Biobank cohort^9^. In the gut, higher *AMY1* dosage has been associated with enrichment of bacteria capable of digesting resistant starch ^8^, potentially supporting more efficient metabolic homeostasis for starch-rich diets. Animal studies have shown that salivary amylase can trigger a cephalic-phase insulin response upon starch consumption ^41^, potentially changing the energy metabolism under starch rich diets. In parallel, numerous studies have attempted to link *AMY1* copy number variation to metabolic traits in humans, with inconsistent results. The largest and best-designed studies to date have not found a direct association with body mass index ^18,42^, suggesting that any metabolic consequences are likely dependent on complex interactions involving diet, environment, and genetic background ^43^.

Within the cultural, dietary, and environmental context of the Andes, the evidence for selection on the amylase locus is concordant with previously suggested starch dietary adaptations among Andeans ^16,17^. Potatoes were one of the first crops domesticated in the Andes, approximately 6,000-10,000 years ago ^11,44,45^. Since then, their descendants have been consuming an extremely starch-rich; a cultural practice that continues today ^13,46^. As mentioned above, researchers have found that the *MGAM* gene, which encodes the maltase-glucoamylase enzyme, shows evidence of selection in Andean populations ^16^ with our samples deriving from a subset of the individuals in Jorgensen et al. This enzyme functions downstream of amylase in the starch digestion pathway, breaking down oligosaccharides generated by amylase activity on complex carbohydrates. Thus, a compelling hypothesis is that the entire starch digestion pathway has been subject to positive selection in Indigenous Andean populations, shaped by a long history of extreme potato-starch intake. Increased *AMY1* copy number, along with complementary changes in genes like *MGAM*, may have jointly optimized starch metabolism to support energy balance and life history traits.

While our findings support adaptation to a starch-rich diet in Andean populations, this hypothesis carries important caveats. The reliance on starchy foods among Andean Peruvians is not vastly different from neighboring groups like the Maya, raising the question of what makes the Andean context unique. One possibility is that a higher *AMY1* copy number may be beneficial for a potato-heavy diet, with potatoes among the highest in glycemic index among all human foods ^12^. However, individuals with higher *AMY1* copy number tend to have increased glycemic responses after consumption of potatoes ^47^. Given that ∼60% of Andean Peruvians carry high *AMY1* copy number haplotypes (≥ 10 copies), the potential for faster starch processing and regulation of glycemia may have conferred an advantage in this setting, but the precise organismal-level effects of this adaptation remain to be tested.

We also observed similarly elevated *AMY1* copy numbers in the Akimel O’odham of Southern Arizona and Northern Mexico. Although sample size precluded formal selection tests, this population presents a distinct ecological and dietary history, rooted in corn-based agriculture and desert environments^48^. If selection on starch metabolism occurred here, it likely reflects different pressures from the Andes, though many further studies are required. Despite their high *AMY1* copy number, the Akimel O’odham, like the Andeans, suffer from high rates of metabolic disease today ^49^, challenging the notion that this genetic variant protects against starch-driven metabolic disorders in modern diets. This pattern aligns with large-scale GWAS studies, which have failed to find strong associations between *AMY1* copy number and obesity-related traits. Interestingly, the strongest biomedical correlate of high *AMY1* copy number is poorer oral health ^9^. This is consistent with the fact that in Peru, over 85% of school-aged children report cavities^50^. While these outcomes are influenced by social and behavioral factors, they highlight the complex health consequences of genetic adaptations shaped by past diets.

By combining long-read sequencing with accurate CNV genotyping, our study offers a framework to study local adaptation at structurally complex loci. Similar approaches could illuminate other multigene families involved in digestion, such as *PGA* and *LPA*. The apparent selection on both *AMY1* and *MGAM* in Andeans suggests potential co-adaptation across the starch digestion pathway. Future work examining how *AMY1* and *MGAM* variants interact to process potato starch will help clarify their specific evolutionary roles. Parallel studies in other high-starch populations, ancient genomes, and functional assays will help test these hypotheses and broaden our understanding of the public health implications of *AMY1* variation globally.

## Supporting information

Supplementary Figures and MethodsTables

## Acknowledgments

We are incredibly grateful to the study participants from Lima, Peru, Cerro de Pasco, Peru, and Palenque, Chiapas, Mexico. We are also thankful to Dr. Maria Inclan (Centro de Investigación y Docencia Económicas), Dr Sudipta Ghosh (NEHU University, Shillong, India), Nate Bartman, Jason Howard, Jacqueline Imse, Kevin Heffernan (Syracuse University), Mark Olfert (University of West Virginia), Francisco Villafuerte (Universidad Peruana Cayetano Heredia), Katarina Evans (CUNY), Neha Angel, Thea Spindel, Sarah Burke, Sarita Greer, Adam Conner, Isabel Meda, and Alber Aqil. This work was supported by NSF grants BIO-2049947 AND BCS-2123284 (to O.G.), BCS-1132310 (to A.W.B., T.D.B., and F.L.-V.) and BCS-1638642 (to A.W.B., T.D.B., and F.S.L.), NIH-R35GM156519 (to O.G.), Leakey Foundation Grant (to A.W.B.), and an Elizabeth Caroline Crosby Grant from the University of Michigan (to A.W.B.).

## Methods

### Sample collection

Peruvian Andean study participants self-identified as Quechua and were recruited from Lima (14 m) and Cerro de Pasco (4,338 m), Peru (**Figure 1E**) as part of a larger study investigating high-altitude adaptation ^21,51^. It is important to note that both Peruvian Quechua study participant groups were of high-altitude indigenous Andean ancestry. Study participants recruited in Lima, Peru, were born, raised, and resided at low altitudes, but whose parents and both sets of grandparents were of highland Quechua ancestry. Peruvian Quechua recruited in Cerro de Pasco, Peru, were born, raised, and resided at high altitudes. Indigenous Mexican Maya participants spoke Tzeltal, Tzotzil, or Ch’ol and were recruited around or in Palenque, Chiapas, Mexico (13 m). Male and female study participants were unrelated, healthy, non-pregnant/lactating, nonsmokers, and between the ages of 18 and 35. All study participants provided whole blood for DNA extraction. Study participants were recruited as part of a larger effort to broadly understand Andean evolutionary adaptation to high altitude. As such, Mexican Maya speakers were selected because 1. they are a close genetic relative of the Quechua who have never lived at high altitude and 2. Previous analysis has demonstrated that Indigenous Americans from Mexico, including Maya, are robust controls for identifying Andean signatures of selection ^31,52^. Importantly for the study of selection at *AMY1*, Andean and Mesoamerican populations diverged prior to the domestication of potatoes ^53^. Furthermore, we were able to sample individuals from both populations (Quechua and Maya) with minimal evidence of recent European or African admixture. All participants provided written informed consent at enrollment for the study. The study was approved by the institutional review boards at the University of California Los Angeles (IRB # 20-001141), the University of Michigan (IRB #HUM00064387), Syracuse University (IRB#22-364), as well as The Research Ethics Committee of the Universidad Peruana Cayetano Heredia, Lima, Peru and the Centro de Investigación y Docencia Económicas, Mexico City, Mexico. Short-read genome FASTQ files from unrelated individuals in the 1000 Genomes Project (PRJEB31736) , the Human genome Diversity Project (HGDP) (PRJEB6463), along with samples from the Simons Genome Diversity Project (SGDP) (PRJEB9586) that belong to populations present in HGDP, were downloaded from the European Nucleotide Archive ^27,28,54^. We additionally utilized high-coverage short-read sequence data from 38 Tibetan genomes ^55^ as well as high-coverage linked read data from 127 individuals from the Middle East ^56^. For all of these datasets, we only kept populations with greater than or equal to eight individuals in order to maximize population diversity and minimize bias due to low sample size.

### Gene Copy Number Calling

To calculate the amylase copy number, we employed two different read depth-based gene copy number callers: mrCaNaVaR and GeneToCN ^57–59^. Both copy number callers have pros and cons. On the one hand, mrCaNaVaR is a promiscuous copy number caller that struggles to differentiate between closely related genes in the same gene families but allows for rapid genome-wide calculations. On the other hand, GeneToCN is quite adept at distinguishing between *AMY1* and *AMY2* genes as well as calculating accurate *AMY1* copy numbers ^3^, but requires much greater involvement in each different gene calculation. To this end, we have opted to use both in our analyses.

For our mrCaNaVaR analyses, using the provided reference sequences and following the reported methods in ^58^ we first aligned each sample FASTQ file to the GRCh38 Human reference genome (CM000663.2) using mrsFAST. We then before calculated the copy number of all genes defined in the reference human_v38.nr.genes.bed file ^60^. We have opted to use the GRCh38 reference genome instead of the newer T2T reference genome because GRCh38 includes three copies of *AMY1* (*AMY1A*, *AMY1B*, and *AMY1C*), which is thought to represent both the ancestral version of the locus and the most common haplotype ^3^, unlike the seven copies present in T2T.

Unlike mrCaNaVaR and most other copy number callers, GeneToCN is an alignment-free method. Instead of calculating read depth across the entire genome, it focuses solely on the read depth of k-mers unique to the *AMY1* genes and conserved flanking regions to determine gene copy number. Following the protocol described in ^59^, we calculated the amylase copy number for *AMY1* genes, *AMY2B*, and *AMY2A* using 25mers.

### VST outlier and median copy number analyses

As the reference human_v38.nr.genes.bed file used in our mrCaNaVaR analysis contains other non-coding genetic features such as long non-coding RNAs, we filtered our dataset for all genes that overlapped with the Ensembl annotation database homo sapiens protein-coding genes from GRCh38 (version 97, ID “AH73881”). We then calculated the VST statistic, VST = (VT−VS)/VT (**see Github**), for all genes that had at least a copy number difference greater than 1 across the dataset in R version 4.4.1 using the protocol described in ^19^ to compare the genes between the populations PEL and MXL along with Quechua samples from Cerro de Pasco and Maya from Palenque. We did not include Quechua samples from Lima in this step due to the differing sequence coverage (∼15X coverage vs. ∼30X coverage). Following this step, we calculated the median of every gene in each population, subtracted the difference between the compared populations, and took the absolute value. After rounding the calculated gene depths and filtering for genes that had a median difference or VST value of 0, we produced a dataset of 576 genes for our PEL and MXL comparison and a dataset of 4,113 genes for our Quechua and Maya comparison. We hypothesized that the difference in coverage between the Quechua and Maya genomes and the PEL and MXL genomes (∼15X coverage vs. ∼30X coverage) might have contributed to a larger number of genes in the Quechua and Maya comparison as fewer reads might have induced greater variance in gene copy number for certain genes that are not in fact copy number variable.

### Admixture analyses

We performed an unsupervised model in ADMIXTURE to determine genome-wide ancestry proportions using 826,848 SNVS after pruning for LD (r^2^ > 0.1) and 1000 Genomes individuals as ancestral references^61^. Individuals were clustered with K=4 into the following super-populations of European, Asian, African, and Indigenous American ancestry. Locus-specific ancestry was assigned using RFMix2 and analyzed via chromosomal segments in windows corresponding to the *AMY* region ^62^. The top 100 1KG individuals from each population with the most corresponding ancestry based on ADMIXTURE results were used as ancestral references for RFMix 2: European (IBS), Asian (CHB/CHS), African (YRI), and Indigenous American (MXL/PEL).

### Quechua & Maya VCF file creation

Whole-genome data for the Quechua samples from Cerro de Pasco and Maya from Palenque, Chiapas, Mexico were generated using an Illumina NovaSeq with 75 base paired-end reads at 15× depth of coverage at the University of Michigan’s DNA Sequencing Core. Whole-genome data for Quechua samples from Lima were sequenced using DIPSEQ-T5 at a higher coverage (∼30X). Low-quality reads with a Phred score < 30 were removed following demultiplexing and trimming with Trim Galore! v.0.6.7 ^63^. Alignment was performed using BWA-MEM against hg38 ^64^, and variant calling was completed with GATK ^65^. Reads were filtered and removed that had low mapping quality, missing per genotype > 0.10, or missing per individual > 0.10 using both VCFtools and Plink2.0 and then phased using SHAPEITv4.2 ^66–68^.

### FST and SNV/*AMY1* association

We filtered out Quechua and Maya individuals with greater than 50% non-American ancestry for the amylase region based on the locus-specific ancestry analysis, removing three Quechua individuals from subsequent analyses (Quechua_Lowland_3, Quechua_Highland_23, & Quechua_Highland_24). Using chromosome 1 data, we filtered for missingness and indels using VCFtools 0.1.16, and retained only biallelic SNVs that had a minor allele frequency greater than 0.05. Following this step, we calculated FST between Quechua and Maya using VCFtools ^69^. Information regarding the frequency of the alternative allele in each population was recorded and used to label if the alternative state was present more frequently in Maya or Quechua samples. For each SNV, we first applied a multiple-hypothesis corrected Kruskal–Wallis test ^70^ (s**ee Github**) **(Table S8 & S9**). To control the family-wise error rate across all SNVs we adjusted the KW p-values with a Bonferroni correction ^71^.

`Kruskal_pvalue <-vapply(snp_data, function(snv) { kruskal.test(AMY1_data ∼ snv)$p.value }, numeric(1))`

We recorded information regarding the median *AMY1* gene copy number for individuals containing homozygous reference, heterozygous, and homozygous alternative states for each SNV and calculated the median gene copy number for each group. Instances where the median *AMY1* copy number of homozygous alternative individuals was greater than either the median of homozygous reference and/or heterozygous individuals, as well as instances where the median *AMY1* copy number of heterozygous individuals was greater than homozygous reference individuals were multiplied by positive one. Instances where the median *AMY1* copy number of homozygous reference individuals was greater than either the median of homozygous alternative and/or heterozygous individuals, as well as instances where the median *AMY1* copy number of heterozygous individuals was greater than homozygous alternative individuals were multiplied by negative one to show a negative association with *AMY1* copy number and the alternative SNV allele.

We calculated a two-sided Wilcoxon test using “stat_compare_means” with comparisons between each SNV allele state (homozygous reference, heterozygous, and homozygous alternative) for two SNVs of interest ^72^ (**Figure 3B & 3D**).

Though several of these SNVs are in and around the *AMY2* genes, none of these SNVs are known to have functional effects on *AMY1* or *AMY2* expression or function and any intronic *AMY2* SNVs that show high LD with selected *AMY1* haplotypes are most likely hitchhiking passengers rather than targets of an independent selection event due to this close proximity.

### XP-EHH Analysis

Using the same filtered SNV file from the FST and Kruskal-Wallis analyses, we calculated XP-EHH between the Quechua and Maya populations using selscan 2.0 ^73^. We created a genetic map file for these SNVs based on human chromosome 1 recombination rates recorded in ^74^. These recombination rates are within the top 95th percentile of chromosome 1. To account for any associations between SNVs on either side of the amylase locus, we extended the haplotype extension to 2 million BP. We additionally applied the Weir and Goudet method in order to account for inbreeding, population structure, or genetic drift that may potentially inflate or distort selection signals ^75^. Finally, we normalized our results across the chromosome. Rs143597860 ranked in the top 23/239,792 SNVs with a XP-EHH value of 6.4.

### Phylogeny

We first filtered the phased SNV dataset for missingness, non-American ancestry at the amylase locus, and to only include biallelic SNVs using VCFtools 0.1.16 ^67^. We then created pseudo haploid FASTA files based on SNVs located between the regions of high recombination surrounding the amylase locus (chr1:103,347,982-103,830,994) for each individual using BCFtools 1.14, SAMtools 1.16.1, and GATK v4.3.0 ^65,76^. To do this, with each individual, we selected only variants where the individual has at least one copy of the alternative state. We then separated these SNVs based on the haploid genome and used the GRCh38 reference genome to create a haploid FASTA file.

`gatk SelectVariants \`

`-R chr1_hg38.fasta \`

`-V${sample}.vcf.gz \`

`--select-type-to-include SNP \`

`--exclude-non-variants true \`

`--O ${sample}_excluded_nonvariants.vcf.gz`

`# For haplotype 1`

`samtools faidx chr1_hg38.fasta ${location} | bcftools consensus -H 1`

`${sample}_excluded_nonvariants.vcf.gz > ${sample}_region_hap1.fasta`

After merging all of the FASTA files, we constructed a maximum likelihood tree by testing multiple models, applying robust branch supports (ultrafast bootstrap), and performing an exhaustive NNI tree search using IQ-TREE 2.2.6 ^77–79^. The output .treefile was labeled with the populations and *AMY1* copy numbers in iTOL version 6 ^80^.

`iqtree -s LWK_Quechua_Maya_no_missing_biallelic.fasta --seqtype DNA -m TESTMERGE -bb 1000 -bnni -allnni -nt AUTO -pre Quechua_Maya_no_missing_biallelic_nni`

Since the phylogeny was constructed using haploid, phased SNV data mapped to the hg38 reference genome, each tip in the tree represents a single haploid genome per individual. However, *AMY1* copy number is measured on a diploid basis. Therefore, individuals with high *AMY1* copy numbers who appear outside the high-copy-number clade in the phylogeny may carry one high-copy haplotype (from the clade) and one low-copy haplotype. This apparent mismatch arises because the phylogeny captures only one haplotype per individual, while the copy number data reflects both. As a result, some individuals with high total copy numbers may possess one high-copy haplotype from the identified clade and another low-copy haplotype present elsewhere in the phylogeny. This interpretation aligns with the data, as 13 out of 14 Quechua SNV haplotypes associated with a diploid *AMY1* copy number of 14, an unusually high count requiring an average of seven copies per haplotype, fall within this subclade.

### Building pangenome

We constructed a local pangenome graph to investigate the mutational mechanisms underlying structural variation at the amylase locus with three specific aims. First, although non-allelic homologous recombination (NAHR) is the dominant mechanism driving *AMY1* copy number variation, other mechanisms such as microhomology-mediated break-induced replication (MMBIR) have been shown to produce structural haplotypes at this locus, including even-copy *AMY1* alleles and *AMY2* variants ^3^. We sought to test whether MMBIR contributed to structural diversity in Peruvian populations. Second, given the recurrent nature of *AMY1* duplications, we aimed to confirm whether previously described recombination breakpoints also underlie the Peruvian haplotypes. Third, we used this graph-based approach to identify potential uncommon haplotypes, including the 7-copy AMY1 haplotype we report here.

The pangenome was built using 40 known haplotypes comprising the entire *AMY* locus ^2,3^. For haplotypes HG02004_pat_H9, HG02293_mat_H5, HG02293_mat_H5, HG02004_mat_H3, HG01978_mat_H3, HG02300_mat_H3, HG02148_pat_H3, HG02293_pat_H1, HG02300_pat_H3, HG01106_mat_H7 ^2^ regions comprising the amy locus were extracted by blasting regions upstream and downstream of the locus. Human reference genome hg38 was used to retrieve the upstream and downstream sequences, respectively chr1:103,547,648-103,560,699 and chr1:103,766,322-103,785,201. For haplotypes H1.1, H1.2, H2A0.1, H2A2B2.1, H3r.1, H3r.2, H3r.3, H3r.4, H3r.5, H3r.6, H3r.7, H3B2.1, H3A2.1, H3A3B3.1, H4A0.1, H4A0.2, H4A2.1, H4A2.2, H4A2B2.1, H5.1, H5.2, H5.3, H5.4, H5.5, H5.6, H5.7, H5.8, H5.9, H5.10, and H7.1, data was retrieved from ^3^) (Supplementary Table 8). All data and scripts used for the pangenome can be found in Github.

We used PGGB ^81^ to build a local pangenome with the following parameters:

`pggb -i amylocus_haplotypes.fa -o amylocus_5k -n 46 -t 32 -p 99 -s 5k`

### Long-read sequencing and assembly

To understand the Peruvian haplotypes for the amylase locus, we chose six Peruvian samples from the 1000 Genomes Project ^28^ to sequence (HG01975, HG01972, HG02106, HG0946, HG01922, HG01955). The samples were chosen to represent the diversity of the population, so we chose samples representing 6, 8, 14, and 18 diploid AMY1 copies, respectively. We generated long-reads on the Oxford Nanopore PromethION sequencing platform using the SQK-LSK114 chemistry and R10.4.1 flow cells for all samples. Because this locus was complicated to assemble correctly, we also generated ultra-long reads for samples HG01975 and HG01946.

We corrected the reads using HERRO ^82^ and assembled the genomes using hifiasm ^83^. We adapted the hifiasm pipeline for the use of corrected ONT reads, as described in ^82^.

`seqkit sliding -s 30000 -W 30000 -g ${cor_reads} > chopped.fasta seqkit seq -m 10000 chopped.fasta > processed.fasta`

`#As ultra-long reads, we are using only reads longer than 50kb seqkit seq -m 50000 $raw > raw_reads_${sample}_50kb.fastq`

`./hifiasm -t 32 -o ${sample}_diploid_50kb --ul raw_reads_${sample}_50kb.fastq -- ul-cut 50000 --dual-scaf -1 parent1.yak -2 parent2.yak processed.fasta`

To verify the assembly recovery of the amylase locus, we aligned the assemblies to the publicly available genome hg38 **(Figure S15**). Based on the dot plots and manual inspection, we concluded that the assembled haplotypes were not correctly assembled. We then used the contigs that span the region to recover the phased reads using the following approach: 1) we mapped all the contigs to the hg38 reference genome; 2) we retrieved the contigs that mapped to the *AMY* locus (coordinates); 3) we retrieved the reads that mapped to the specific parts of the contigs that mapped to the region of interest.

All scripts used for this are on **Github**.

### Mapping to haplotypes and filtering

Because the assembled contigs were incorrect, we mapped the reads to all known distinct haplotypes (n=30) plus previously published Peruvian haplotypes ^2^ and used a read depth measure to decide the best fit for each sample. The haplotypes used are on **Github**, details in **Table S15**. We competitively mapped the reads into each haplotype and filtered the mapping for length > 2.5kb. This filtering was done because the region is rich in transposable elements that might skew the analysis.

We hypothesized that the best-fit haplotype will have a lower variance in mapping. Therefore, we calculated the mean and standard deviation for the read depth on the entire haplotype. The best fit for all haplotypes was the one that showed the lowest standard deviation and a mean read depth of ≥ 1. We also checked that the best-fit haplotype for each individual coincided with the estimated copy number based on ddPCR results (**Table S13**). Based on this method, we could assign the best fit for 4 individuals (8 phased haplotypes).

However, two individuals (HG02106 and HG01975) were assigned haplotypes discordant with the estimated copy number generated by ddPCR. We manually inspected the best-fit haplotypes, and the mapping was not homogeneous for any of the top haplotypes (H9, H7, H5). The best match for this haplotype was H9, and, therefore, we hypothesized that the best match would be a haplotype similar to H9 but harboring 7 *AMY1* copies. We generated a mock H7 haplotype by deleting one duplication unit (*AMY1A* and *AMY1B*, see ^3^) from the H9 haplotype (**Supplementary Methods, Figure S16**).

### Data availability

Results generated for this study are provided as Supplementary Data 1-15. Additional datasets are provided in this paper including the PBS results, the Ohana results, the XP-EHH results, the mock H7 FASTA file, and the SNV LD analyses have been uploaded to Github https://github.com/kendrasc/Andean_Amylase_Project. Phased SNV data between the locations of Chr1: 103343000-103960000 in the Quechua and Maya individuals is available on Figshare along with archived versions of the aforementioned results and datasets where they are freely available without restriction. The 1KG data and the HGDP data can be obtained at https://www.internationalgenome.org/data/. The SGDP data can be obtained at https://www.simonsfoundation.org/simons-genome-diversity-project/. The raw sequencing data generated for this study have been deposited in the NCBI Sequence Read Archive (SRA) under BioProject accession number PRJNA1240630. The FASTQ files are available under the following SRA accession numbers: HG02106 (SRR32828259), HG01946 (SRR32828263), HG02252 (SRR32828258), HG01975 (SRR32828261), HG01972 (SRR32828257), HG01922 (SRR32828264), HG02006 (SRR32828260), HG01955 (SRR32828262). Each sample is associated with a BioSample record (accessions SAMN47540692 to SAMN47540699) and represents Homo sapiens individuals of Peruvian ancestry from the 1000 Genomes Project.

For peer review, the data can be accessed via the following private link: https://dataview.ncbi.nlm.nih.gov/object/PRJNA1240630?reviewer=2reo2tk31pprot2497moukd0e2 The whole genome data that support the findings of this study may be available upon reasonable request to the corresponding authors.

### Code availability

The code used to generate the results in this study is publicly accessible on GitHub https://github.com/kendrasc/Andean_Amylase_Project.

