## Supplementary Figures and MethodsTables for "Rapid Adaptive Increase of Amylase Gene Copy Number in Indigenous Andeans"

**Adaptive Increase of Amylase Gene Copy Number in Peruvians Driven by Potato-rich Diets**

**The file includes:**

Supplementary Results

Supplementary Materials and Methods

Figs. S1 to S16

References

**Supplementary Methods** 3
1.1 Additional Gene Copy Number Testing 3
1.2 VST calculations 3
1.3 LD between AMY1 Copy Number and SNVs across Populations 3
1.4 Calculating Quechua and Maya population histories and effective population sizes 4
1.5 Simulating SNV frequencies over time in Quechua and Maya populations to test for probability of rs143597860 real-life frequencies. 4
1.6 PBS Analysis 5
1.7 Ohana Analysis 6
1.8 Assigning best-fit haplotypes for long-read data and generating a mock-H7 haplotype 6
1.9 Evaluating mutational mechanisms 7
**Supplementary Figures:** 9
Figure S1. Boxplot and Kruskal-Wallis test and p-values for various geographic regions 9
Figure S2. Boxplot of AMY2A copy numbers across the world 10
Figure S3. Boxplot of AMY2B copy numbers across the world 11
Figure S4. The association between AMY1 copy number and flanking SNVs in the Quechua, Maya, and 1000 Genomes populations for all SNVs that pass a Bonferroni correction of 0.05 12
Figure S5. r2 across the amylase and flanking regions as associated with rs1930184 and rs143597860 14
Figure S6. Phylogeny of Quechua, Maya, and EAS populations for SNVs flanking the Amylase locus. 16
Figure S7. Quechua and Maya population history 17
Figure S8. Fraction of SNVs that had a frequency of ≥ 45% in Quechua and ≤ 13% in Maya individuals (the same percentages as rs143597860) based on backwards simulations of SNVs out of Africa for different effective population sizes. 18
Figure S9. XP-EHH across chromosome 1 19
Figure S10. rs143597860 for PBS, XP-EHH, and Ohana tests 20
Figure S11. TMRCA of SNV rs143597860 using a joint clock model. 21
Figure S12. CLUEs results for rs143597860 in Maya and Quechua that include posterior probability values. 22
Figure S13. CLUEs result for EGLN1 SNV rs1769792 23
Figure S14. Illustration of how non-allelic homologous recombination (NAHR) can generate different haplotypes at the Amylase locus. 24
Figure S15. Alignment of phased contigs to the amylase locus on the reference genome hg38. 25
Figure S16. Generation of a mock H7 haplotype. 26

### Supplementary Methods

#### 1.1 Additional Gene Copy Number Testing

Using the *AMY1* copy numbers calculated by GeneToCN [1,2] for all individuals, we performed a Kruskal-Wallis test on each designated geographic region using ggpubr, "stat_compare_means" (**Figure S1**) [3].

#### 1.2 $V_{ST}$ calculations

The biggest challenges to calculating $V_{ST}$ across all protein-coding genes were results that were influenced by genome coverage with smaller coverages resulting in greater gene copy number variances that would affect the calculated copy number, mrCaNaVaR's inability to distinguish between closely related genes, and potential variation caused by partial gene duplications such as VNTRs instead of entire gene duplications. Though we could not look into every gene directly, we checked and filtered out four genes that were calculated to have over 100 copies in the genome and that had an average median copy number difference between the populations far greater than all other genes. Though we removed these genes from our analysis, we do note their differences between the Quechua and Maya populations. These genes were:

1. **USP17L family** → This gene family ranges from 20 to 103 copies, due to the RS447 mega satellite region that has been shown to change from generation to generation (~8.3% change in transmission) [4]. Due to the high frequency of gene copy number turnover, we chose to remove this gene family from our analyses.
2. **NBPF family** → This gene family generally has around 26 copies but an Olduvai repeat domain within the genes that can range within the hundreds seems to be driving the high population population differentiation as opposed to the actual gene copy number variation. We are not looking into this type of copy number variation, so this family is removed [5].
3. **MTRNR2L8** → BLATing this gene's sequence in the HG38 reference genome shows that this gene is a pseudogene caused by a NUMT that seems to have inserted itself all over the genome [6].
4. **DUX4** → This gene can range to over 100 copies and like the USP17L gene family, is caused by the D4Z4 macrosatellite near the end of chromosome 4 and has the largest range of copies of any ORF in the human genome [7]. As we are not looking at this type of copy number variation (caused by microsatellites), this gene was removed.

#### 1.3 LD between *AMY1* Copy Number and SNVs across Populations

We then ran Kruskal-Wallis tests with a Bonferroni correction between flanking SNVs and *AMY1* copy number on each population in the Quechua, Maya, and 1000 Genomes project (**Figure S4, Table S8 & S9**). The most significant values belonged to SNVs in the Quechua population. Interestingly enough, EAS populations additionally had a large number of SNVs positively significantly associated with *AMY1* copy number. Most of these SNVs have a shared significance across the EAS and American populations, potentially hinting that the high copy number haplotypes driving the high population average copy number in Andean populations are also responsible for the high population average copy numbers in East Asians as mentioned by earlier work [8,9]. Additionally, there is high LD between SNVs shared between EAS and American populations in the Quechua population (**Figure S5**). A phylogeny of the SNVs flanking the amylase locus in Quechua, Maya, and EAS populations shows that individuals with high copy number haplotypes in Americans fall in a higher-than-average *AMY1* copy number clade shared between East Asian and American populations (**Figure S6**).

#### 1.4 Calculating Quechua and Maya population histories and effective population sizes

We used *Relate* to estimate the effective population size history and divergence times for the Quechua and Maya populations [10]. We first filtered our Quechua and Maya chromosome 1 phased SNV dataset for missingness before removing the three individuals that had amounts of non-American ancestry in the amylase locus. To run relate, we used the provided "genetic_map_chr1.txt" for our genetic map along with the default mutation rate of 1.25e-8, an N of 30,000, and a generation rate of 28. We first ran "RelateFileFormats --mode ConvertFromVcf" before running "Relate --mode All", followed by EstimatePopulationSize.sh with bins of 3, 7, and 0.2 as used in the example.

#### 1.5 Simulating SNV frequencies over time in Quechua and Maya populations to test for probability of rs143597860 real-life frequencies.

To test the probability of a SNV reaching the frequency of rs143597860 in Quechua and Maya populations, we did backward simulations using Msprime [11,12]. Using frequencies provided in gnomAD and Atlas of Variant Age modern populations rs143597860 is present in the Americas (15% in 1000 Genomes, 31.8% in Simons Genome Diversity Project, and 24% in HGDP) with Peruvians and Pima having the highest frequencies in the datasets [13,14]. The only presence in other populations is 3.7% in Central Asian Siberians, 0.333% in Europeans from HGDP, and 0.1% in Africans (although there is a discrepancy as this number on gnomAD is 0%). In other much larger datasets rs143597860 is present 174/67356 in Non-Finnish Europeans (0.2583%), 74/41484 in Africans (0.1784%), and 1/4792 in South Asians (0.02087%). Since this SNV has a low frequency and does not appear at all in East Asian populations that are the closest relatives to Americans, we do not know if this is a very old SNV that has been segregating at low frequencies for thousands of years, or if these patterns are the result of convergent evolution. As the current data indicates that before entering the Americas, the SNV would have been at a very low frequency that only increased after the bottle neck into the Americas, rather than simulate SNVs throughout the history of humans, we chose to begin our simulation with Out of Africa.

In order to obtain dates and effective population sizes, for the Quechua and Maya populations during and after their split, we ran Relate [10] (**see methods 1.3**). Because our data becomes less reliable further back in time—potentially producing artifacts like a population decline around 100,000 years ago instead of a later decline consistent with other Out-of-Africa studies—we rely on effective population sizes and split times from other research for earlier historical events in our analyses [15–18]. As these numbers are often ranges, we have opted to take the averages.

Ancestral effective population size: 13,600

| | | |
|---|---|---|
| Out of Africa: | Time = 60,000 years ago* | Ne = 1,750** |
| Americas Bottle Neck | Time = 16,000 years ago*** | Ne = 2,125 |
| Quechua/Maya Split | Time = 10,000 years ago**** | Ne prior = 3,000 |

Time = 6,000 years ago Ne Maya drop = 2500

Time = 1,000 years ago Ne Maya end = 3200

Time = 1,000 years ago Ne Quechua end = 1500

*We set the time of Out of Africa at 60,000 years ago rather than the average of 52,500 years ago to be more conservative and allow more time for SNVs to appear in the population
**While [18] estimates a higher effective population size for populations leaving Africa (> 2,000 individuals), we have opted to use the lower estimate of 1,750 from [16] in our simulations. This lower effective population size enhances the role of genetic drift during bottlenecks, making our simulation results more conservative in terms of drift effects.
***The crossing of the Bering land bridge had several stages with the divergence between Siberians and proto-Native Americans occurring 24,900 years ago, the settling of Alaska with the Native American founder lineages occurring 18,400 years ago, and the expansion into the rest of the Americas 16,000 years ago. We have chosen to use the date at 16,000 years ago as this bottle neck is representative of the populations that eventually became the Maya and the Quechua.
****We used a split time of 10,000 years ago, based on our own calculated data, instead of the North/South American split time of 14,775 years ago cited in [19], to account for potential interactions following the initial continental population separation.

This simulation was run for 3,000,000,000 bases (around the size of the human genome) with a generation time of 25 years and a mutation rate of 1.165e-8. We then re-ran this simulation 192 times to test the effects of lower effective population sizes. We then calculated the fraction of SNVs that had a frequency of $\geq 0.45$ in Quechua and $\leq 0.13$ in Maya.

#### 1.6 PBS Analysis

We calculated PBS using SNVs from Chromosome 1 in the FIN, PEL, and MXL 1000 Genomes populations. We filtered the files to keep only biallelic SNVs that had a minor allele frequency greater than 0.05 resulting in a total of 5,013,617 SNVs. This file is larger than the VCF files created from the Quechua and Maya data due to known information regarding allele status of SNVs present in populations outside of the Americas. Following this step, we calculated pairwise $F_{ST}$ between the populations using VCFtools 0.1.16 [20,21]. We then ran PBS on each in r using the $F_{ST}$ results based on the instructions found on https://mccoy-lab.github.io/hgv_modules/index.html. Rs143597860 ranked in the top 2,027 SNVs with a PBS value of 0.31.

#### 1.7 Ohana Analysis

We performed a population-specific selection scan on chromosome 1 using Ohana (v0.1.0; https://github.com/jade-cheng/ohana) [22]. Phased VCF files for Quechua individuals were merged with 1000 Genomes Project reference populations IBS (Iberian from Spain) and YRI (Yoruba from Ibadan, Nigeria) using bcftools-merge (v1.17) [23], retaining only biallelic SNPs. IBS and YRI were selected as reference populations based on prior unsupervised Admixture results. Variants with missing genotypes were removed in PLINK (v1.90b6.26) [24], and the filtered dataset was converted to Ohana's genotype format (.dgm) using ped2dgm.

A random subset of ~5% of SNPs was selected with sample-sites.py and used to estimate ancestry proportions (Q matrix) and allele frequencies (F matrix) with qpas (K=3). A covariance matrix (C) was generated from the subset using nemeco and modified to create a contrast-specific matrix (cs.matrix) by adding a constant to the diagonal entry corresponding to the Quechua component.

The full dataset was then analyzed with selscan using the fixed Q matrix from the subset (-qi) and the contrast-specific C matrix (-cs). SNPs with likelihood ratio test statistics above the 99th percentile were considered candidates for selection. The full analysis script is available at: https://github.com/kendrasc/Andean_Amylase_Project/blob/030b1ef00363047dc282743a8721aca6fc5086a7/Supplementary/Figure_S10/ohana_run.sh. Rs143597860 ranked in the top 69 SNVs.

#### 1.8 Assigning best-fit haplotypes for long-read data and generating a mock-H7 haplotype

We initially aimed to assemble the amylase haplotypes using both long-read and ultra-long-read data generated by Oxford Nanopore sequencing. Assembly statistics are provided in **Table S8**. However, due to the extreme complexity and high repeat content of this locus, we were unable to generate a reliable assembly. We tested multiple strategies, including: (1) hifiasm [25] with ultra-long reads and error-corrected ONT reads, (2) Verkko2 [26] with trio-based data (incorporating short-read sequences from the parents of 1000 Genomes individuals), and (3) local assemblies using Flye [27], hifiasm, and Canu [28]. None of these approaches produced a complete and accurate reconstruction of the locus; the best results consisted of either collapsed assemblies or fragmented contigs that failed to span the full amylase region (**Figure S16**). Furthermore, the inferred *AMY1* copy number from these assemblies did not match the expected values determined by ddPCR.

This method, however, was useful to extract the phased reads for each contig, given hifiasm generates phased contigs with coordinates for which reads made up for each contig. When raw reads are input into hifiasm, the assembler uses heterozygosity patterns at specific sites within the reads mapping to a given region to phase them, producing two distinct haplotypes. Hifiasm also outputs a mapping of each contig to the reads used to assemble it, allowing retrieval of all reads contributing to a specific region. We first aligned the phased contigs to the hg38 reference genome and extracted those mapping to the amylase locus (hg38 chr1:103,570,000–103,760,000). We then retrieved all reads associated with each contig and realigned them to the contig. Finally, we isolated only the reads mapping to the amylase locus (including reads that may start earlier or end later than the locus), thereby obtaining phased reads for the amylase locus for each individual. The pipeline for extraction of phased reads is in **Github**.

We then decided to map the phased reads to the known haplotypes, in order to define the best match for each phased sample. We mapped all phased samples to all haplotypes previously reported by [29] (H1.1, H1.2, H2A0.1, H2A2B2.1, H3.1, H3.2, H3.3, H3.4, H3.5, H3.6, H3.7, H3B2.1, H3A2.1, H3A3B3.1, H4A0.1, H4A0.2, H4A2.1, H4A2.2, H4A2B2.1, H5.1, H5.2, H5.3, H5.4, H5.5, H5.6, H5.7, H5.8, H5.9, H5.10, H7.1). Additionally, we included all the Peruvian haplotypes alongside with the longest reported haplotype, H9, reported by [9] (HG01978_mat, HG01978_pat, HG02004_mat, HG02004_pat, HG02148_pat, Hg02293_mat, HG02293_pat, HG02300_mat, HG02300_pat, HG01106_mat, with 3,5,3,9,3,5,1,3,3,7 *AMY1* copies respectively). This totalized 40 haplotypes.

The best haplotype match for each sample should have more evenly distributed mapped reads throughout the entire haplotype. With this premise, we calculated the mean and standard deviation for the read depth for each mapping (**Table S10**). Two criteria made up for our choice of best match sample-haplotype: the mapping with the lowest standard deviation and at least one read covering each region of the haplotype (which translates to the mean read depth being $\geq 1$). To further validate this, we compared the results of the best-match haplotypes with the ddPCR results, and we kept only the samples that were consistent across the two methods. With this, we were able to estimate which haplotypes are present in the Peruvians sequenced using the long reads for four of the six individuals sequenced.

When carefully inspecting the mappings, we noticed that the best matching for the phased samples HG02106 and HG01975 was to H9 haplotype, and this did not match with the expected by ddPCR results (14 copies diploid). We then manually inspected the mapping and saw a dip in mapping that spanned two repetitions of the *AMY1* gene (the region in grey in Figure S1). By manually deleting that region, we were able to generate a mock H7 haplotype (**Figure S1**) from the H9 haplotype. We then mapped both phased samples to this new haplotype. It is of note that we kept the architecture of the locus as described by [29], meaning we deleted not only the genes, but the DNA blocks that make up the intergenic regions between the genes, according to **Figure S1**, aiming to more accurately map the reads. We successfully assign the best mapping for both phased samples to the H7 mock haplotype, which is in accord with the ddPCR results.

#### 1.9 Evaluating mutational mechanisms

We evaluated the mutational mechanisms underlying copy number variation observed in the amylase loci for nine individuals sequenced specifically for this study (HG01946, HG01922, HG01972, HG01975, HG02106, HG01955, HG02252, HG02006, and HG02106). To determine whether any of the 18 assembled amylase loci harbored non-recurrent, single-origin MMBIR haplotypes, we utilized the previously characterized MMBIR stretches described in [29]. Three haplotypes have been previously defined as complex, non-recurrent rearrangements with microhomologies at their breakpoint junctions: H2A2B2.1, $H3^r$.6, and H3B2.1. We isolated the MMBIR microhomology stretches and extended them by 75 bp upstream and 75 bp downstream from the identified breakpoint junctions. These extended sequences were queried using BLASTN to identify matches against the whole genome of the ancestral GRCh38 reference haplotype $H3^r$.2, serving as a negative control, and the whole genome haploid assembly of the individual carrying the MMBIR haplotypes, serving as a positive control [30]. This analysis verified the presence of a single perfect hit for each MMBIR breakpoint junction stretch in the genome assembly harboring the specific MMBIR haplotype. In contrast, no perfect or continuous hits were recovered when searching against a genome assembly without the MMBIR haplotype. None of the 18 assemblies produced hits for any of the queried MMBIR stretches. However, it is important to note that additional complex, non-recurrent, single-origin MMBIR haplotypes may exist that have not yet been identified and characterized, and therefore could not be accounted for in this analysis. Nonetheless, based on the currently resolved structural haplotypes, none of the 18 assemblies, and by extension the nine sequenced individuals, provide any evidence supporting the presence of MMBIR amylase haplotypes. The queried MMBIR stretches are provided below:

>H3$^r$.6_MMBIR_stretch
AAATATTTAAACAATGTTTTGCATTTCTCTGATGGCCAGTGATGATGAGCATTTTTTCATGTG
TCTTTTGGCACATGAAACCCTGTCTCTACTAAAAATACAAAACAAATTAGCCAGGCATGGTG
GTGGGTGCCTGTAGTCCCAGCTACTCA
>H3B2.1_MMBIR_stretch
GAAGCATGGAGAAGTGAACCCTGGAGCTTCCCAATGGTGAGATACCAAATAGTTCAAGAAC
TAGTAAAGAAGACTACTTAAAAATAAGAGCTAGGCACAGGGATTAAAATATATACTTCGAA
TAAGTACCTACCTCAGGGTTAATAGGAAGA
>H2A2B2.1_MMBIR_stretch

GGTATGGGCCATTCTCCTGCCTCAGCCTCCTGAGTAGCTGGGACTACAGGCACCCACCACCA
TGCCTGGCTAATTTGTTGTTAGTGGTTATTTTATTTTTATATTTATCAACAATATCTTCATTTC
CTAATTGAGAACCTAGTATGAAATCCAT

### Supplementary Figures:

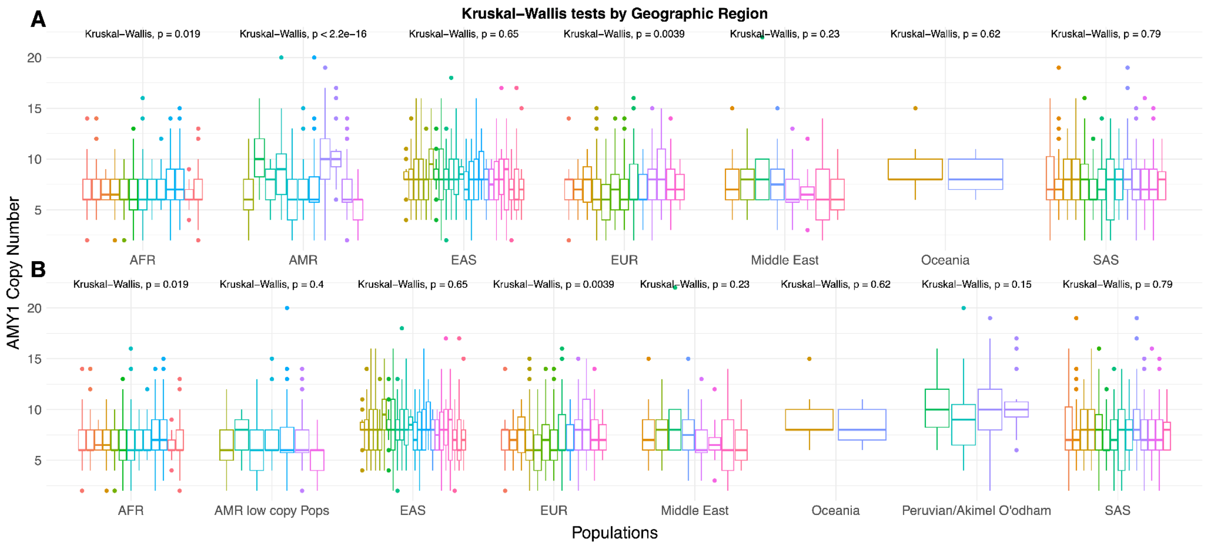


**Figure S1. Boxplot and Kruskal-Wallis test and p-values for various geographic regions**

Boxplots of *AMY1* copy number calculated using GeneToCN for all samples tested in this dataset. Kruksal-Wallis p-values have been calculated for each population. We note that the p-value is significant for populations in the Americas when the Peruvian and Pima populations are included (**Figure S1A**), but not when they are missing (**Figure S1B**). Additionally, the Peruvian and Pima populations do not have significant differences from each other.

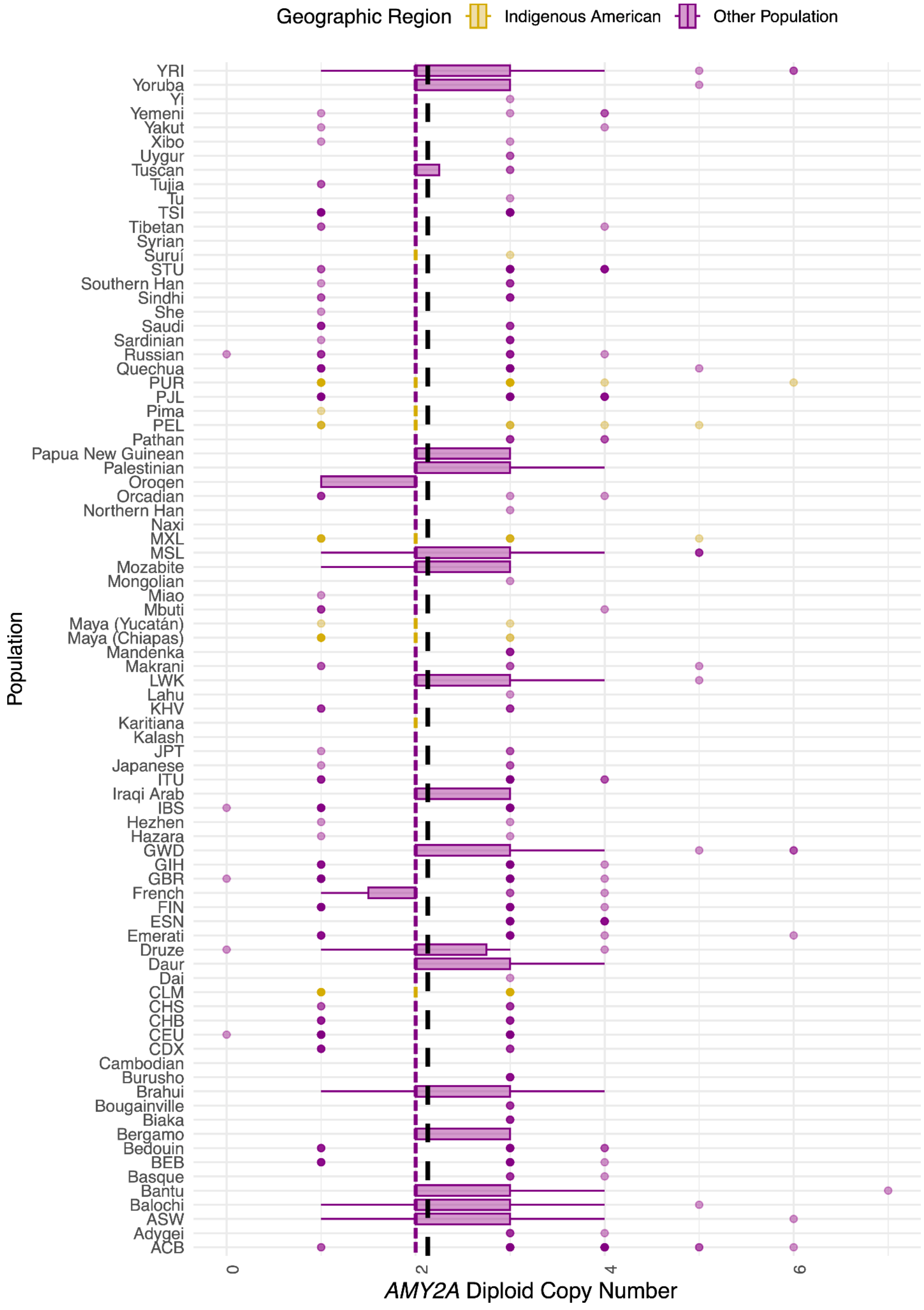


**Figure S2. Boxplot of *AMY2A* copy numbers across the world**

Populations are compiled using samples from the 1000 Genomes dataset, the HGDP dataset, individuals from Tibet and the Middle East, as well as Quechua and Maya individuals we have provided. Gold represents American populations. Purple represents non-American populations. Worldwide mean is labeled in a dotted black line. There is no increase in *AMY2A* copy number in Peruvian populations as compared to other American populations.

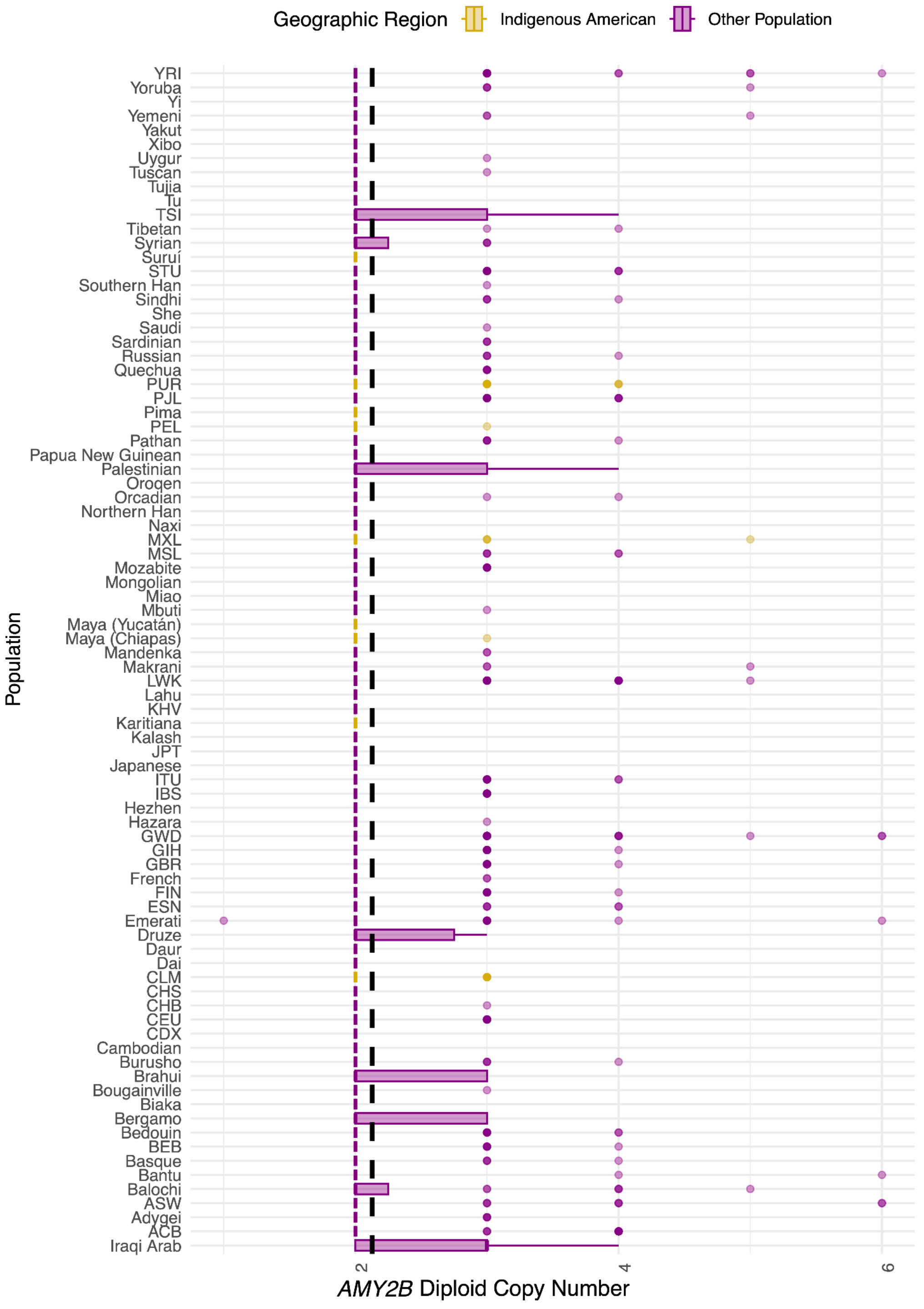


**Figure S3. Boxplot of *AMY2B* copy numbers across the world**

Populations are compiled using samples from the 1000 Genomes dataset, the HGDP dataset, individuals from Tibet and the Middle East, as well as Quechua and Maya individuals we have provided. Gold represents American populations. Purple represents non-American populations. Worldwide mean is labeled in a dotted black line. There is no increase in *AMY2B* copy number in Peruvian populations as compared to other American populations.

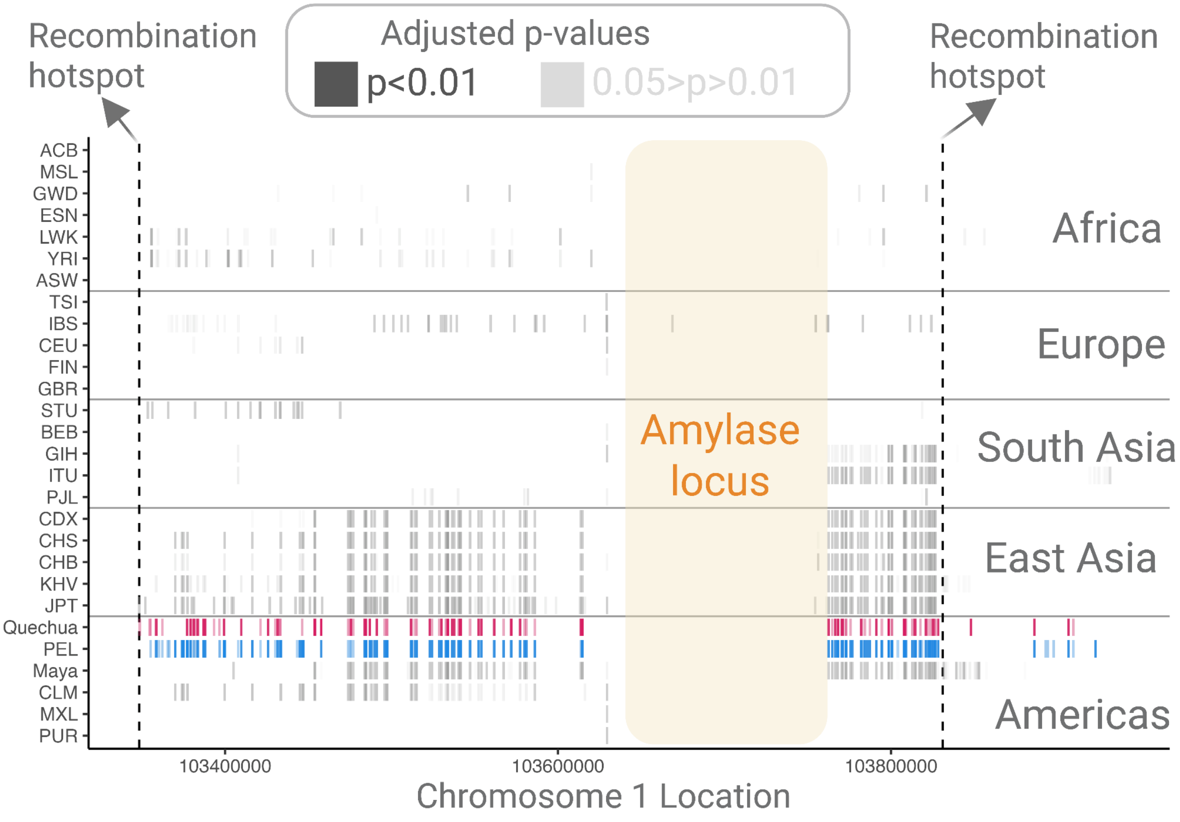


**Figure S4. The association between *AMY1* copy number and flanking SNVs in the Quechua, Maya, and 1000 Genomes populations for all SNVs that pass a Bonferroni correction of 0.05**

The x axis shows a SNVs location on chromosome 1 around the amylase locus with the regions of high recombination labeled, while the y axis splits the SNVs up by population (**Table S8**). Only SNVs that are significantly associated with *AMY1* copy number under a Kruskal-Wallis test and pass a Bonferroni correction of 0.05 in a population are labeled. The darker the shading, the more significant the p-value is. The Quechua and PEL populations are also colored in red and blue for ease of viewing purposes.

Most populations do not have any, or have few SNVs significantly associated with *AMY1* copy number including European populations. This result aligns for the most part with the previous results. For example, in [31], though there does seem to be some relationship between flanking SNVs and *AMY1* copy number shared across European populations, the $r^2$ results between SNVs and *AMY1* copy is quite low.

The main outliers in this dataset are East Asian populations and un-admixed American populations.

East Asian populations all have high amounts of SNVs associated with *AMY1*. For reference, in CDX (Chinese Dai) which has the least amount of significantly associated SNVs, there are still 81/228 total SNPs (~ 36%) in the flanks that are significantly associated with AMY1 copy number after adjusting for Bonferroni corrections. On the other end of the spectrum, JPT (Japanese in Tokyo) which has the highest number of significantly associated SNVs has 135/227 total SNPs (~59%) in the flanks significantly associated with *AMY1* copy number after adjusting for Bonferroni corrections. All 81 SNVs significant in CDX are also significant in CHS (Han Chinese in the South), CHB (Han Chinese in Beijing), JPT, and

KHV (Kinh from Ho Chi Minh City, Vietnam). Most of these SNPs seem to be within the regions of high recombination on either side of the locus.

The Quechua, Maya, PEL, and CLM (Colombian from Medellín) populations display high numbers of associations between *AMY1* copy number and flanking SNVs. Notably, PEL and CLM have the highest proportions of Indigenous American ancestry among the 1000 Genomes American populations [32]. Additionally, the same 81 SNVs shared across all East Asian populations are also significant in the PEL and Quechua samples, 77 of which are also significant in Maya (**Table S9**).

These results hint at a shared haplotype across East Asian and American populations. This haplotype appears to be breaking up in the Maya population, as the number of significant SNVs decreases with distance from the amylase locus, especially compared to the Andean populations. A similar pattern is observed in the CLM population, where significant SNVs are only found upstream of the locus. While several SNVs remain significant downstream of the amylase locus in Maya, beyond the region of high recombination, closer inspection suggests these may tag low-copy-number haplotypes, potentially indicating a loss of the *AMY1A* and *AMY1B* block that may stifle recombination in the region.

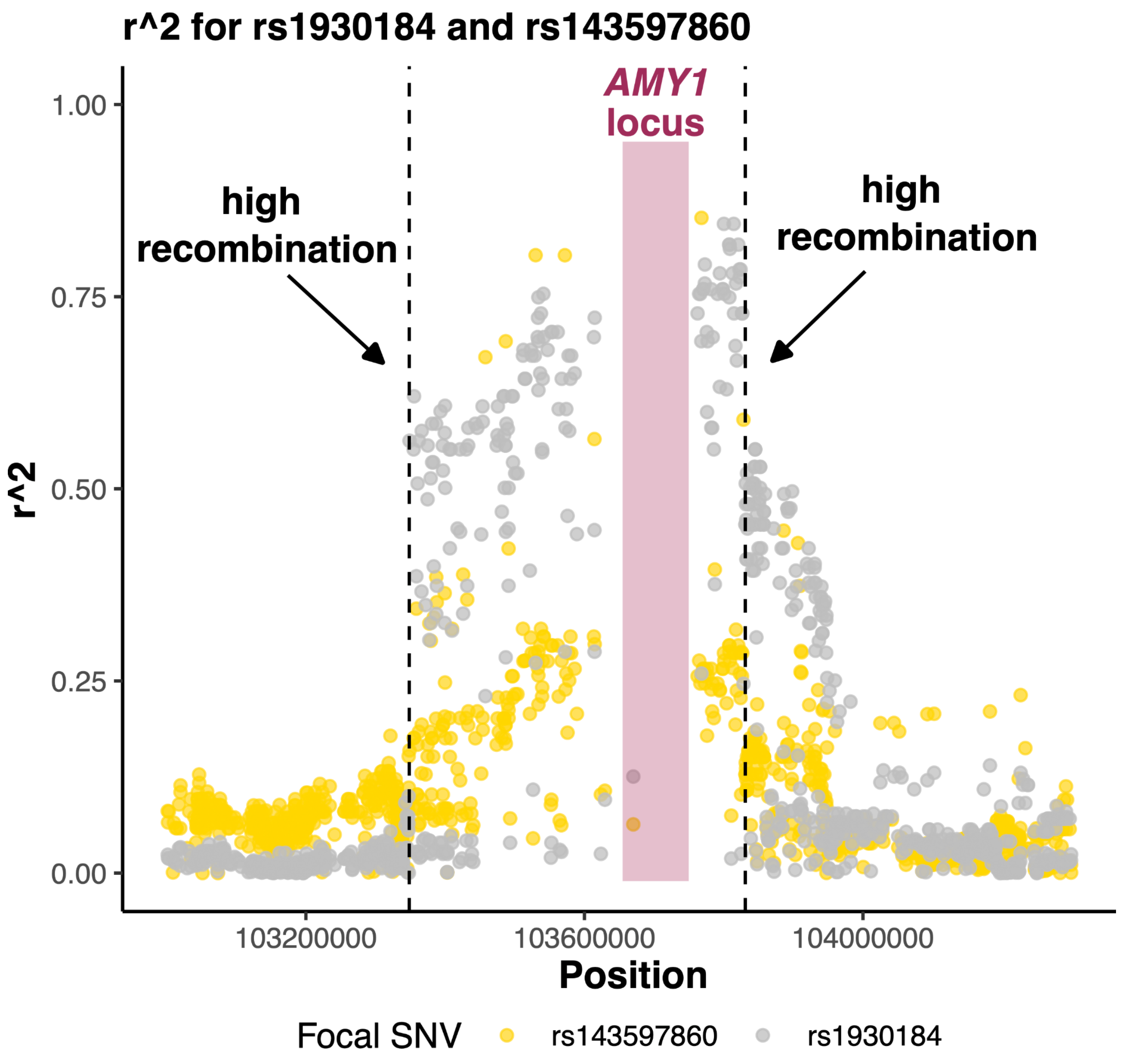


**Figure S5. $r^2$ across the amylase and flanking regions as associated with rs1930184 and rs143597860**

We calculated $r^2$ values using VCFtools for SNVs between 103,000,000 and 104,300,000 in the Quechua population. This calculation only included SNVs that are biallelic and have a minor frequency greater than 0.05. The regions of high recombination flanking the amylase locus are labeled with a black dashed line and the amylase locus itself is highlighted in burgundy. We then filtered to include only $r^2$ values between SNVs and rs1930184 (labeled in grey) or rs143597860 (labeled in gold).

In **Figure 3**, we note that rs1930184 tags a cluster of SNVs in Quechua and Maya (cluster 1) where the reference allele is associated with higher *AMY1* copy numbers and is a high frequency in Quechua (~74%) and a lower frequency in Maya (~42%). This SNV is also one of the 81 SNVs from **Figure S4** that are associated with high AMY1 copy number in East Asians. Also, in **Figure 3A**, we highlight rs143597860 as a SNV tagging a cluster of SNVs (cluster 3) that have the alternate allele associated with high *AMY1* copy number and segregating at a higher frequency in Quechua (~46%) than Maya (~14%). We note that rs143597860 has not been found in a single individual of East Asian in all public datasets and is segregating at low frequencies (< 1%) in other populations where it is present.

Overall, SNVs directly flanking the amylase locus within the regions of high recombination tend to have high $r^2$ values with rs1930184. This aligns with the results from **Figure S4** that indicates that there is a shared ancestral haplotype between East Asians and American populations that is responsible for the high associations between *AMY1* copy number and flanking SNVs in those populations. Rs143597860, on the other hand, despite being the most significantly associated SNV with *AMY1* copy number across the 1000 Genomes project in addition to the Quechua and Maya populations, tends to have lower $r^2$ values with the flanking SNVs excluding several other SNVs that are also associated with high *AMY1* copy number in the alternate state. Due to this trend, we hypothesize that rs143597860 at some point after the split between the ancestors of modern day East Asians and the ancestors of modern day Americans joined a pre-existing SNV haplotype that was already highly associated with *AMY1* copy number. Though the version of this haplotype carrying rs143597860 rose in frequency in the Andean population to be carried by almost half of the population, versions of this haplotype lacking rs143597860 are still present, resulting in imperfect linkage disequilibrium.

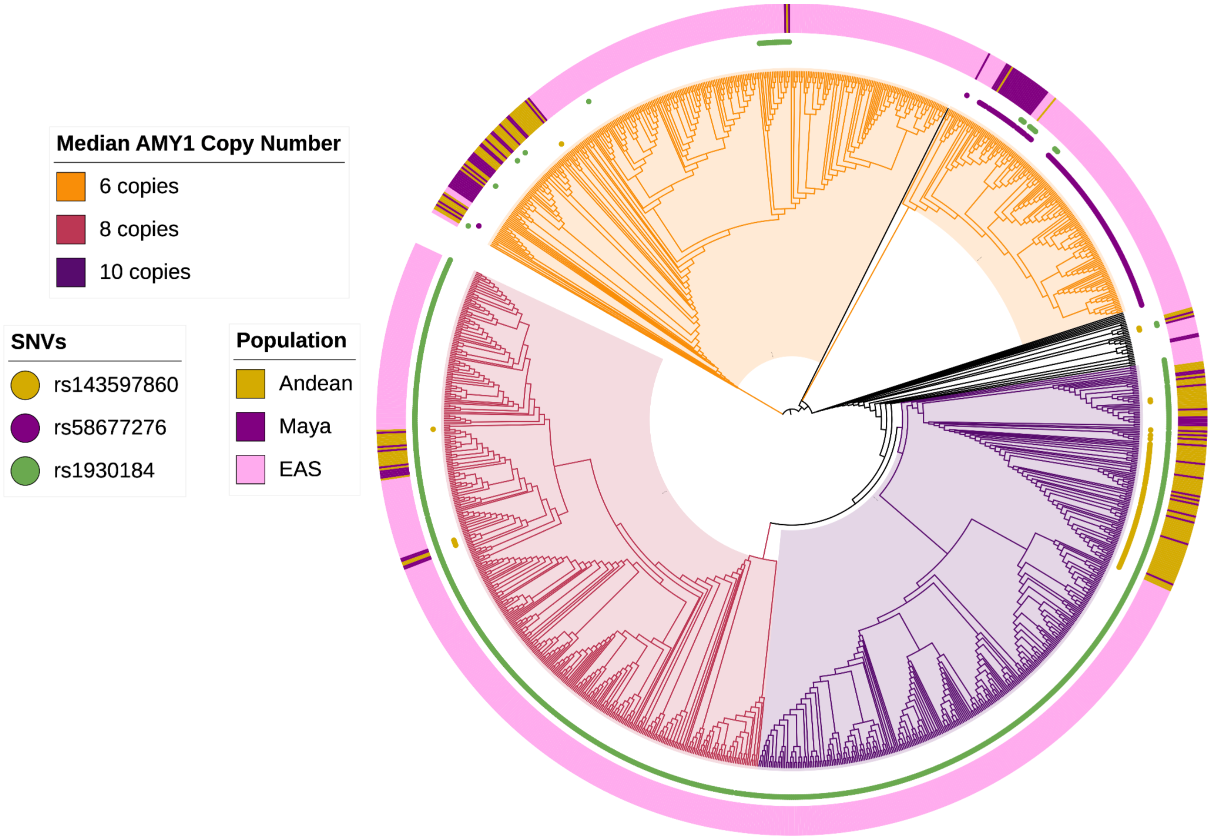


**Figure S6. Phylogeny of Quechua, Maya, and EAS populations for SNVs flanking the Amylase locus.**

The **Figure S6** phylogeny is made of the amylase locus and its flanking regions up to (HG38 Chr1: 103,348,464-103,830,994) using phased VCF files and the HG38 reference genome. Ascertainment bias for invariant sites controlled for using the command "--ASC" in IQ-TREE.

The first ring of color labels the population each individual belongs to (EAS, Maya, or Quechua). The subsequent circles label the presence or absence of SNVs tagging specific clades. Green circles denote the presence of the rs1930184 reference allele which tags a large high copy number (Median $\geq 8$) clade. Purple highlights the presence of rs58677276 which tags the low copy number samples in Maya (**Figure 4C**). Gold circles indicate the presence of rs143597860, the SNV that tags the high copy number clade in Andeans.

The branches are labeled based on the median copy number of the clade they belong to (if they belong to a large clade). Orange branches have a median of 6 diploid *AMY1* copies, burgundy branches have a median of 8, and dark purple branches have a median of 10. The high copy number American clade that has expanded in Andeans is nested in the dark purple clade, indicating that these high copy number haplotypes had already existed, and that this potential selection event was on standing variation on a haplotype that may have been present in the ancestors of East Asians and Americans, giving further evidence to the hypothesis mentioned in **Figure S5**.

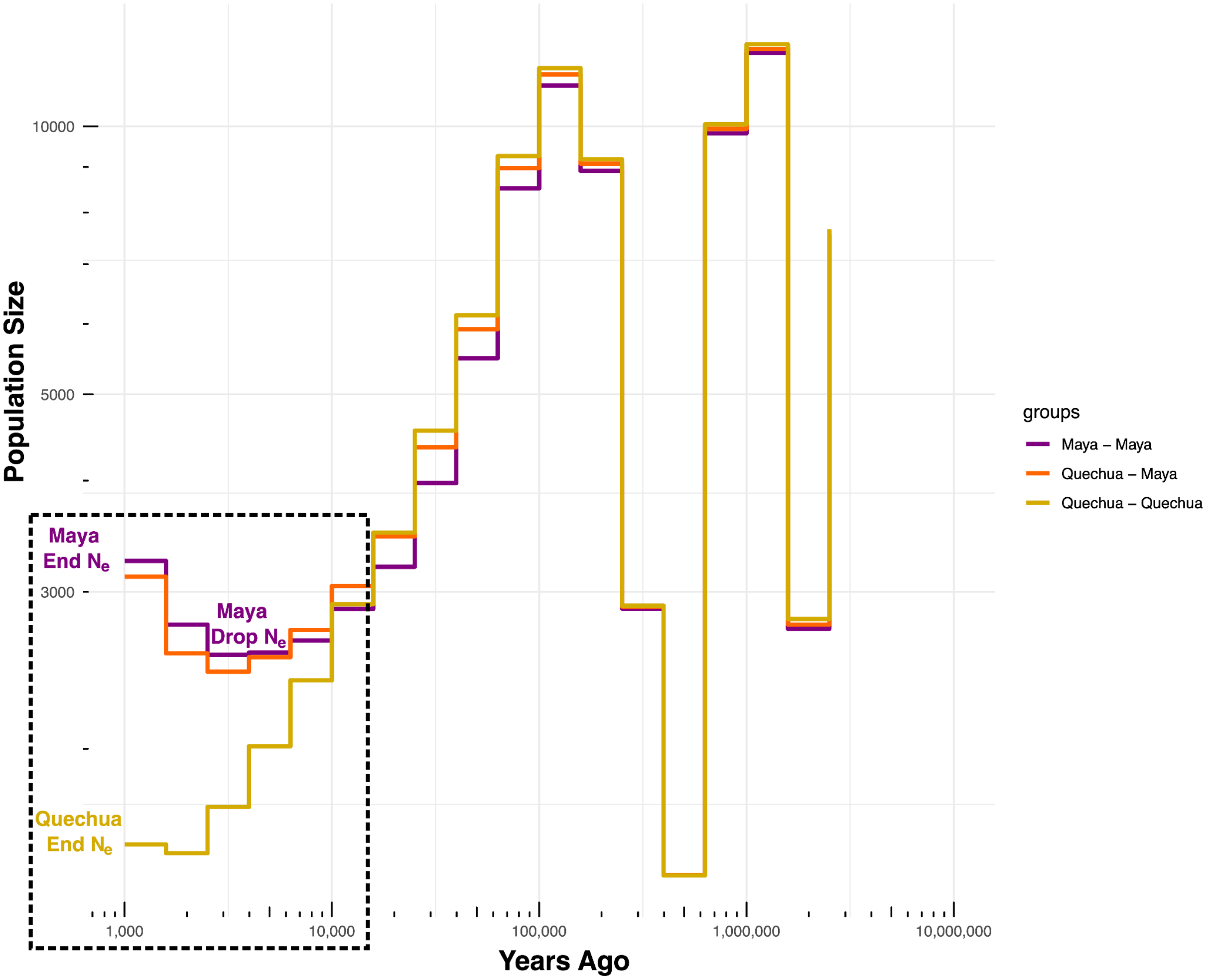


**Figure S7. Quechua and Maya population history**

We modeled the population history of Quechua and Maya individuals using phased SNV data in Relate. This model informed key events and effective population sizes, including a population split around 10,000 years ago and the final effective population sizes for Maya and Quechua individuals in our simulated SNV datasets. Sizes and dates that are used are highlighted in the dotted black rectangle.

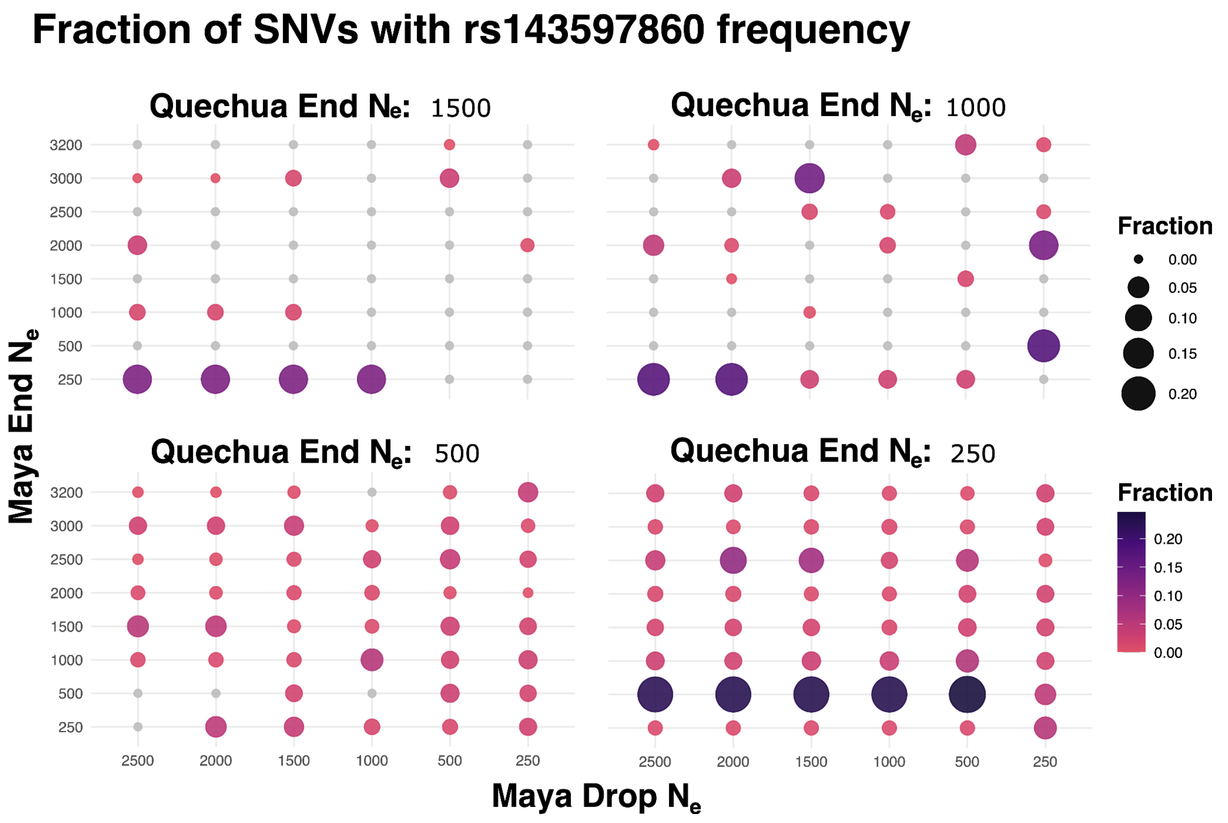


**Figure S8. Fraction of SNVs that had a frequency of ≥ 45% in Quechua and ≤ 13% in Maya individuals (the same percentages as rs143597860) based on backwards simulations of SNVs out of Africa for different effective population sizes.**

The top right corner point effective population sizes (Quechua End Ne = 1500, Maya End Ne = 3200, and Maya Drop Ne = 2500) are derived from **Figure S8**. The end effective population size of the Quechua population had the greatest impact on whether or not these frequencies were reached. All simulations run at an end Quechua Ne of 250 produced SNVs that appear at these frequencies while other effective populations decreased in likelihood the greater the effective population size was. The ending Maya Ne additionally had an effect, especially if the end Maya effective population size was 500 people and Quechua was 250 people.

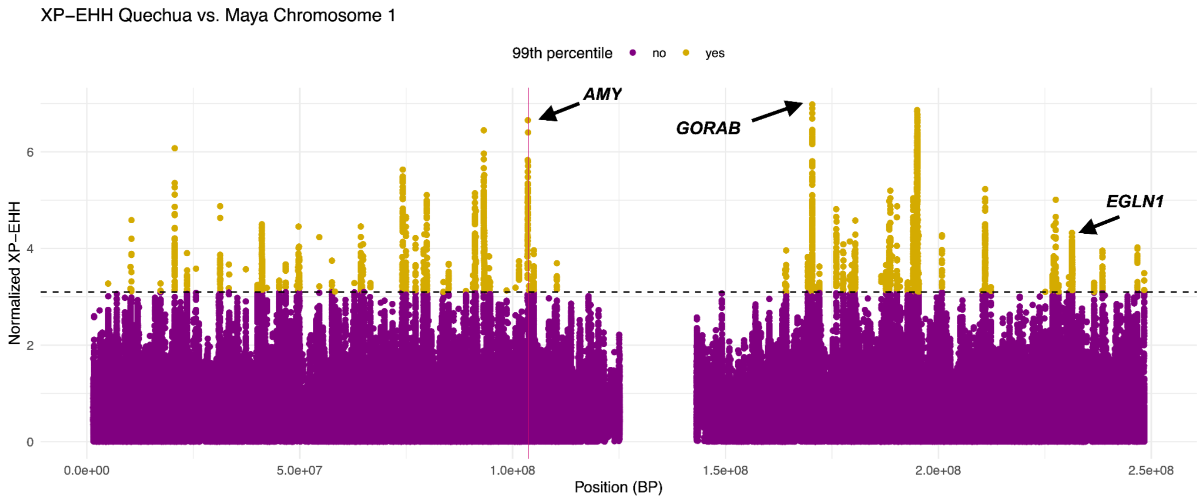


**Figure S9. XP-EHH across chromosome 1**

XP-EHH across chromosome 1 with *AMY*, *GORAB* (the top gene), and *EGLN1* (a gene involved in high altitude adaptation [33,34] highlighted. The top SNV associated with GORAB increases its expression in the arteries according to GTEx data [35].

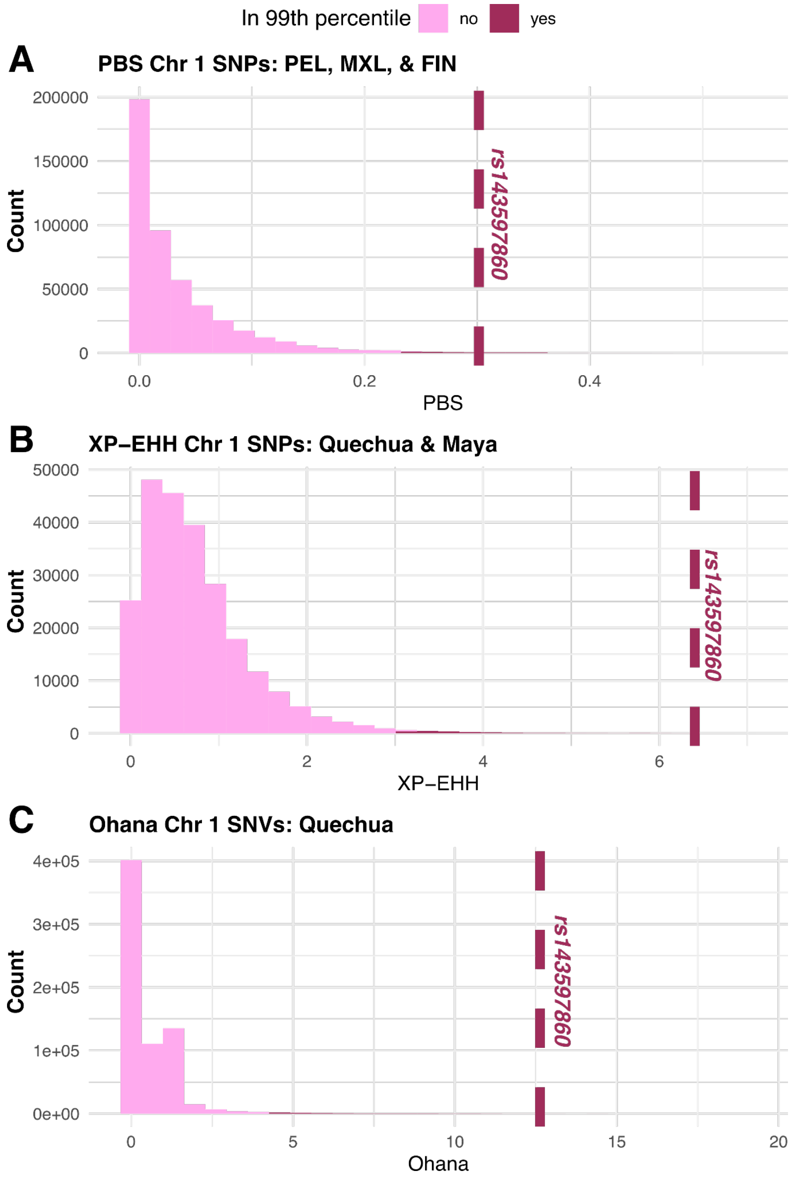


**Figure S10. rs143597860 for PBS, XP-EHH, and Ohana tests**

Three selection tests on chromosome 1 highlighting rs143597860. Values in burgundy are in the 99$^{th}$ percentile. The value of rs143597860 is detonated by a dashed line in each figure. We applied the Population Branch Statistic (PBS) using $F_{ST}$ calculated using VFtools and followed a tutorial from the Human Genome Variation Lab led by Dr. Rajiv McCoy and published on the lab's official site (https://mccoy-lab.github.io/hgv_modules) [36]. This was calculated using the PEL, MXL, and Finnish in Finland (FIN) populations from the 1000 genomes dataset. The XP-EHH results are the same results as in **Figure 4A** but expanded to include the entire chromosome. We ran Ohana on the Quechua population with Iberians in Spain (IBS) and Yoruba in Ibadan, Nigeria (YRI) as comparisons to take into consideration potential admixture from the Iberian Peninsula and western Africa. In each case, the calculated value for rs143597860 is within the 99$^{th}$ percentile, indicating that it is an outlier.

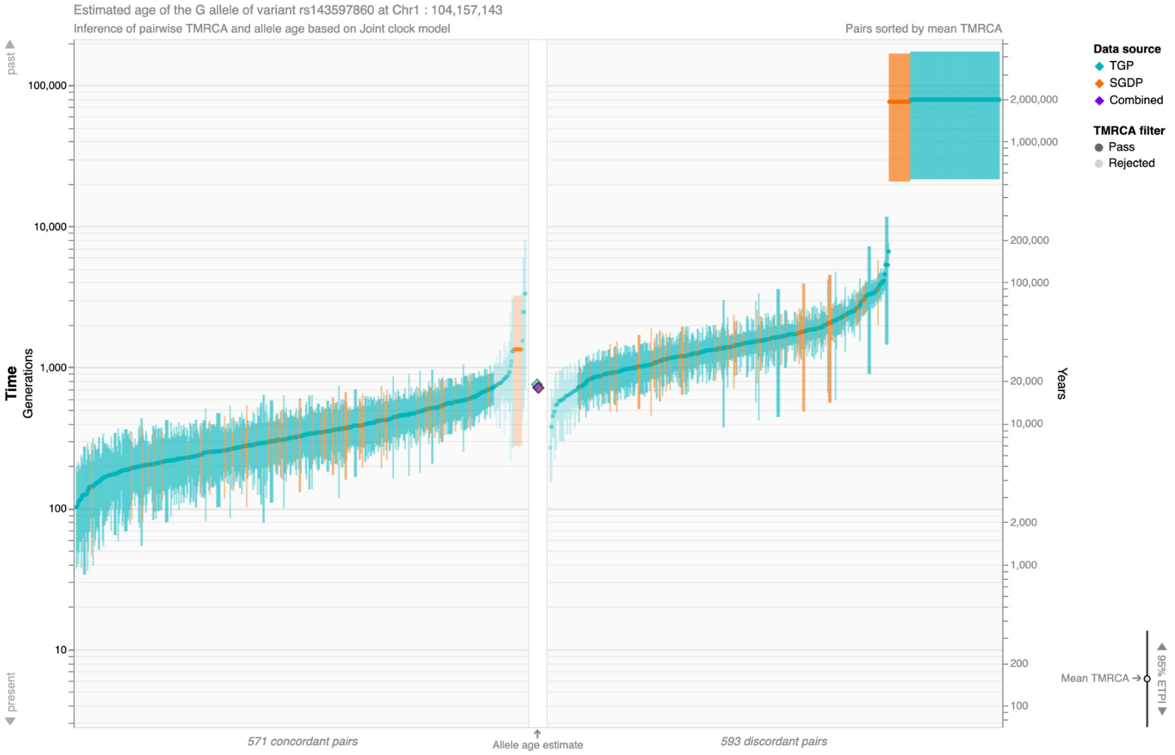


**Figure S11. TMRCA of SNV rs143597860 using a joint clock model.**

We utilized the Human Genome Dating: Atlas of Variant Age database to estimate the age of rs143597860 [13] Dating of the allele is based on coalescent times of concordant and discordant pairs on phased genomes using Genealogical Estimation of Variant Age (GEVA). Based on the 1000 Genomes (TGP) and Simons Genome Diversity Project (SGDP) dataset, the alternative allele for rs143597860 is dated a little before the crossing of the Bering Strait, with the SNV segregating at a frequency of 3.7% in populations from Central Asia Siberia before rising to 15-31% in the Americas. Samples in blue are estimates using samples in the TGP dataset while samples in orange are from the SGDP dataset. These results are concordant with the estimated trajectory of this SNV based on the CLUEs analysis which estimates low frequencies before a relatively recent increase in the last 10,000 years (**Figure 4B, S12**).

**A.**

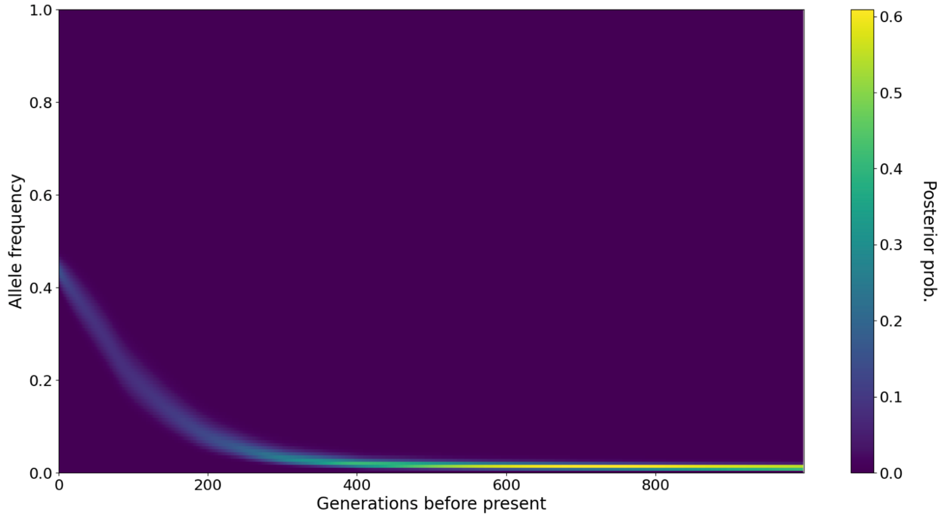


**B.**

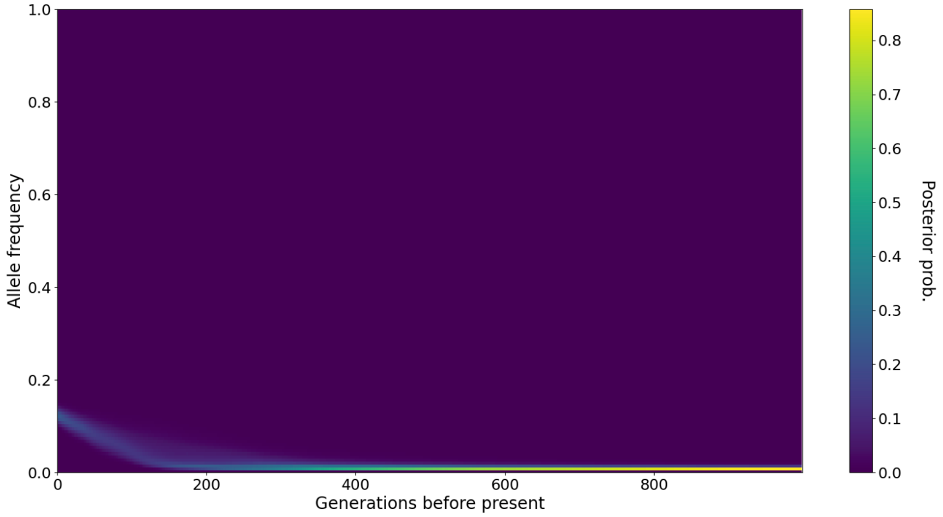


**Figure S12. CLUEs results for rs143597860 in Maya and Quechua that include posterior probability values.**

**Figure 12A** shows the estimated trajectory for rs143597860 in Quechua while **Figure 12B** shows the trajectory in Maya. Results were calculated using Relate and CLUES. The posterior probability decreases more recently in time, especially around the timing of the increase in frequency of this SNV in each population.

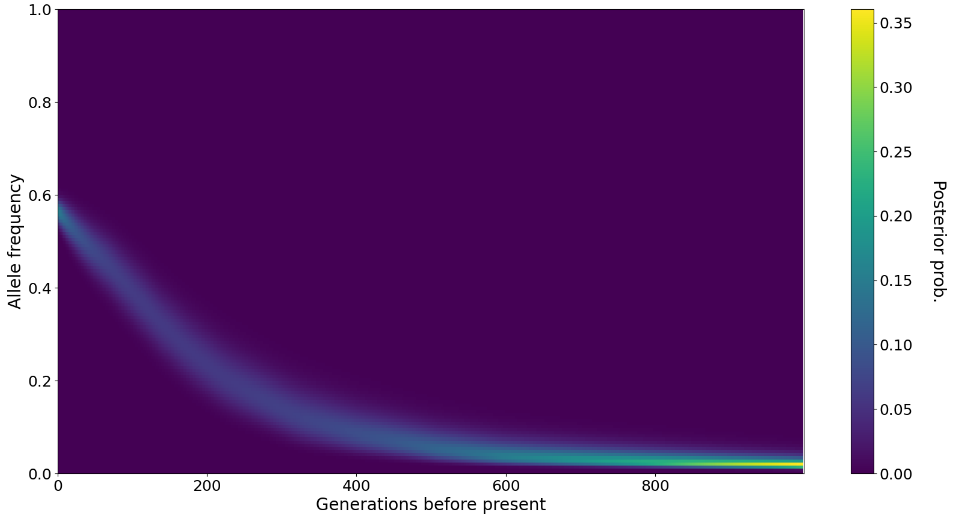


**Figure S13. CLUEs result for *EGLN1* SNV rs1769792**

Estimated historical trajectory for SNV rs1769792 Andean populations based on modern day Quechua individuals. Results were calculated using Relate and CLUES. The posterior probability decreases more recently in time, especially around the timing of the increase in frequency of this SNV in each population.

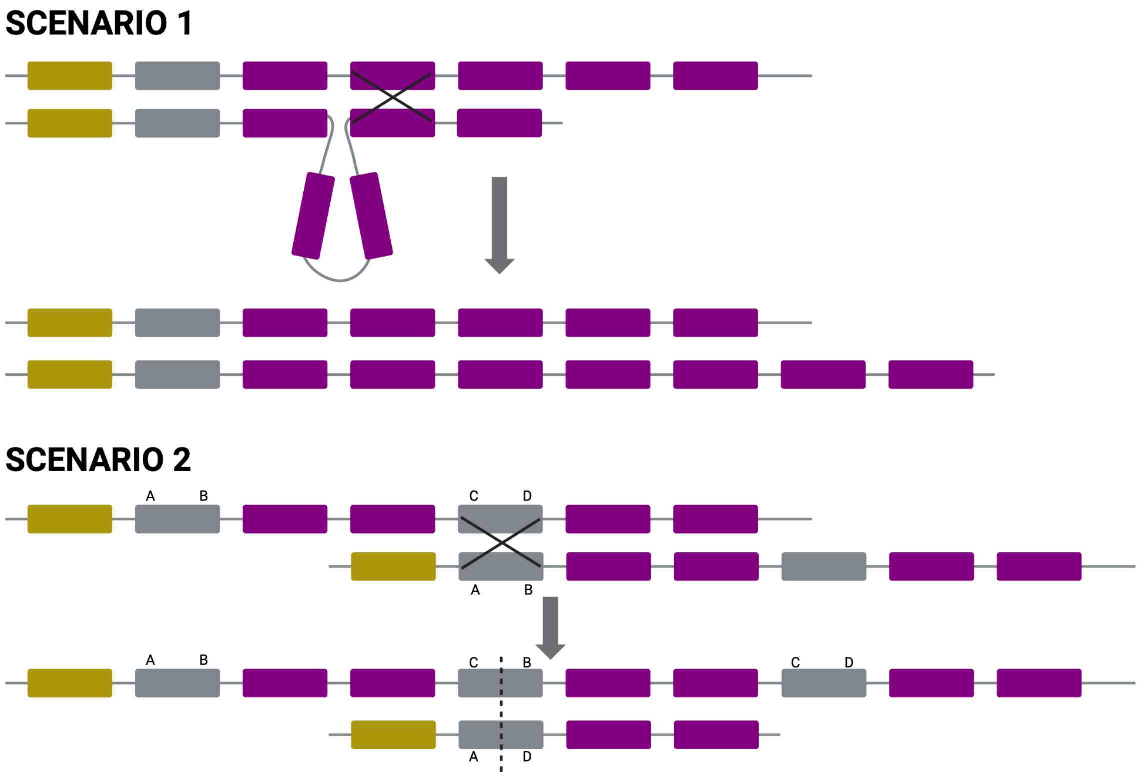


**Figure S14. Illustration of how non-allelic homologous recombination (NAHR) can generate different haplotypes at the *Amylase* locus.**

In both scenarios, colored segments represent hypothetical homologous DNA sequences found in common haplotypes. In Scenario 1, a loop forms during homologous pairing in recombination. A segment of DNA pairs with a downstream homologous segment, and the replication machinery incorporates an extra copy into the resulting haplotype. This results in a haplotype containing an additional homologous segment. In Scenario 2, a grey segment pairs with a downstream grey segment, and recombination occurs within the middle of the segment. As replication proceeds along the paired strand, this event produces one haplotype with a duplication of the grey segment and another with a deletion of the same segment.

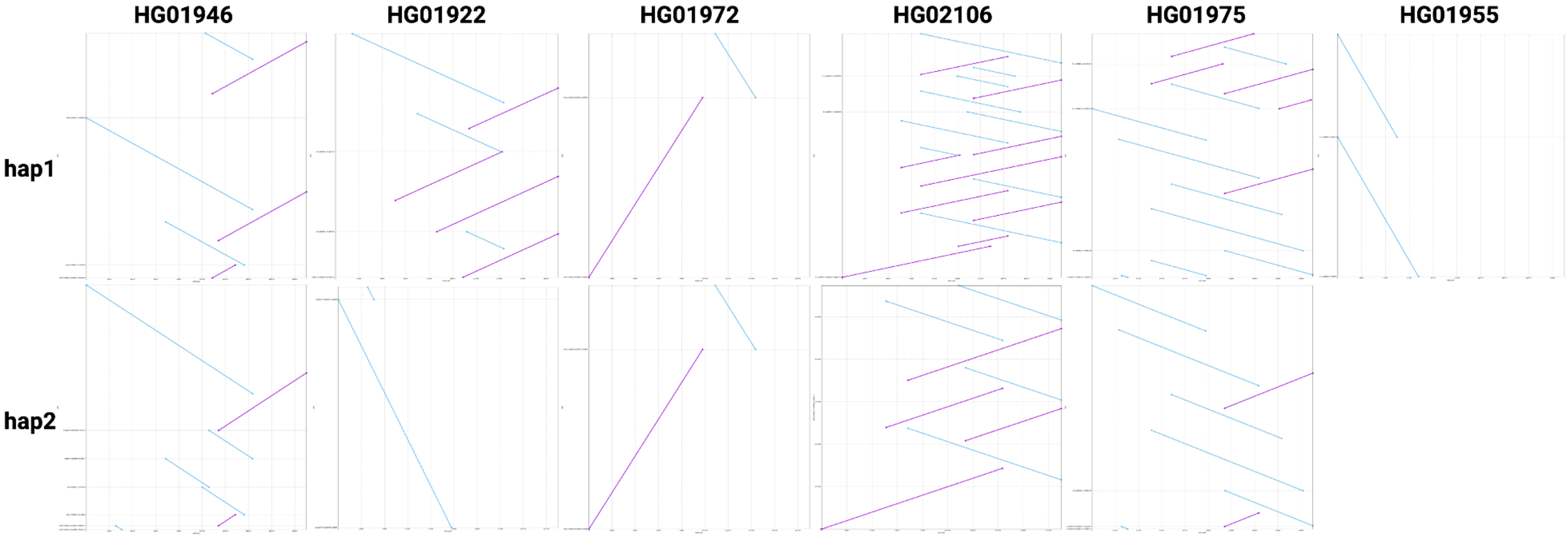


**Figure S15. Alignment of phased contigs to the amylase locus on the reference genome hg38.**

On the x axis, the *AMY* locus (hg38 chr1:103570000-103760000), and on the y axis, the contigs that map to that region. The sample HG01955 had no contig mapping to the region on haplotype.

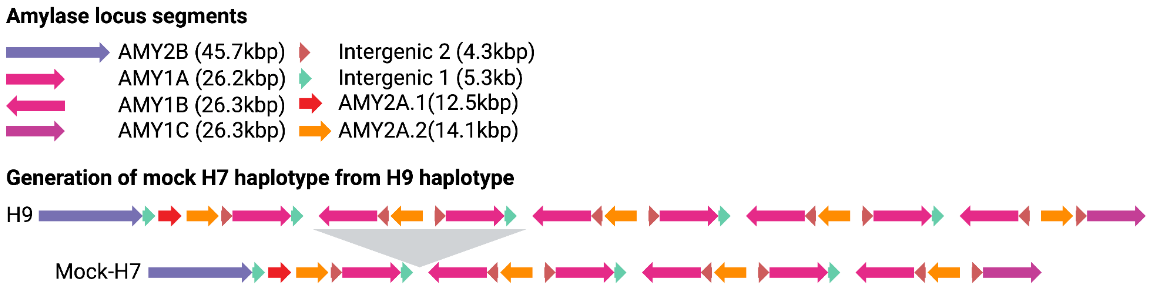


**Figure S16. Generation of a mock H7 haplotype.**

On the top, we see the description of the DNA segments that make up the Amylase locus, as described by [29]. At the bottom, a schematic representation of the region of the H9 haplotype that was deleted to generate an H7 mock haplotype. The DNA sequence of the mock H7 haplotype can be found in (https://github.com/kendrasc/Andean_Amylase_Project/tree/main/Figure_5

36. *Human Genome Variation Lab*.
